# Systems-level properties of EGFR/Ras/ERK signalling amplify local signals to generate gene expression plasticity

**DOI:** 10.1101/466656

**Authors:** Alexander E. Davies, Taryn E. Gillies, Stefan Siebert, Michael Pargett, Savannah J. Tobin, Abhineet R. Ram, Celina Juliano, Gerald Quon, Mina J. Bissell, John G. Albeck

**Affiliations:** Department of Molecular and Cellular Biology, University of California, Davis, CA; Division of Biological Systems and Engineering. Lawrence Berkeley National Laboratory, Berkeley, CA

## Abstract

The EGFR/Ras/ERK signalling pathway is a driver of cancer cell proliferation and metastasis in tumours that exhibit high cell-to-cell heterogeneity. While the signalling activity of this pathway is frequently amplified in tumours, it is not understood how the kinetic aspects of its activation in tumours differ from normal cellular signalling. Using live-cell reporters of ERK signalling in the breast cancer progression series HMT-3522, we found that ERK activity in invasive cells is similar in amplitude to isogenic non-malignant cells but is highly dynamic and more disordered, leading to more heterogeneous expression of ERK target genes. Our analysis reveals that this diversification arises from systems-level functions of the pathway, including intracellular amplification of amphiregulin-mediated paracrine signalling and differential kinetic filtering by genes including Fra-1, c-Myc, and Egr1. Our findings establish a mechanism for the generation of non-genetic tumour cell plasticity arising from the specific quantitative properties of a signal transduction pathway.

## Introduction

Heterogeneity at multiple levels is a prominent feature of many solid tumours, including aggressive basal-like breast cancer (BLBC)^1, 2^. Whereas some heterogeneity can be attributed to the genetic mosaicism of tumours, it is now clear that much of the cell-to-cell variation arises from cellular plasticity, defined as the ability of tumour cells to reversibly shift their gene expression profile and phenotypic characteristics over time^3^. Plasticity provides an adaptive advantage for cancer cells that contributes to metastasis and drug resistance. Potential drivers of plasticity are multifactorial^4^ and include both intrinsic stochastic processes and extrinsic signalling and adhesive inputs received from the tumour microenvironment (TME)^5, 6^. However, it remains unclear how these factors are translated into cellular heterogeneity and whether they are sufficient to drive plasticity.

The Ras/ERK pathway is an important candidate for driving heterogeneity, as it controls many characteristics of invasive cancer cells and has been observed to vary widely from cell-to-cell in culture and *in vivo*^7, 8^. Genes stimulated by this pathway include transcription factors such as c-Myc and Fra-1 that have been implicated as drivers of malignant behavior in breast cancer^9-11^. While BLBC tumours do not frequently carry mutations in the Ras cascade, they often overexpress epidermal growth factor receptor (EGFR)^12^ or display protein phosphorylation profiles consistent with receptor tyrosine kinase activity^13^. This suggests that even in the absence of mutations in the pathway, extracellular cues in the tumour microenvironment (TME), including TGF-α, amphiregulin (Areg), and certain extracellular matrix (ECM) molecules can promote Ras activation^14^. Recent advances in genetically encoded reporters now enable dissection of transient, short-range Ras/ERK signalling events generated by the microenvironment^15, 16^. These experiments have revealed that Ras/ERK signalling displays a high degree of dynamic heterogeneity that is modulated by growth factors, other autocrine/paracrine signalling, and cell density^17^. These observations suggest the hypothesis that dynamic Ras/ERK signalling is closely linked to the plasticity of cells and their ability to shift their gene expression profile frequently.

In this study, we investigated the link between Ras/ERK signalling dynamics and gene expression plasticity associated with BLBC using the HMT-3522 progression cell line model^18^. These cells were originally derived from a reduction mammoplasty and subjected to multiple rounds of *in vitro* and *in vivo* selection for tumourigenic behavior^19^. This culture model allows for the study of cellular properties at each stage of spontaneous, rather than genetically engineered, tumour progression. Using cells from different stages expressing live-cell ERK reporters, we identify a dynamic Ras/ERK signalling profile present only in malignant cells. We show that this dynamic ERK originates in paracrine signalling through amphiregulin that is then amplified by the EGFR/Ras/ERK signaling pathway, driving continual fluctuations in gene expression in adjacent malignant cells and co-cultured non-malignant cells. These signals result in stochastic time-varying expression of cancer-related genes, establishing microenvironment-mediated Ras/ERK signalling dynamics as a mechanism sufficient to drive cellular gene expression plasticity.

## Results

### Spontaneous Ras/ERK signalling dynamics are altered in malignant progression

To investigate the relative differences in Ras/ERK signalling dynamics in non-malignant and malignant cells, we generated HMT-3522 cells carrying genetically encoded fluorescent reporters of ERK activity (Figure 1A). Non-malignant (S1) and malignant (T4-2) cells were transduced with the ERK translocation reporter (ERKTR), and we measured ERK activity by live-cell microscopy as the ratio of cytosolic (C) to nuclear (N) fluorescence (hereafter referred to as ERKTR_C/N_). We evaluated S1 and T4-2 cells under: 1) baseline imaging medium lacking exogenous EGF and serum, or other MAPK activators, 2) treatment with MEK inhibitor, PD0325901 (PD) or 3) stimulation with EGF. At baseline conditions, S1 cells exhibited few fluctuations in ERKTR_C/N_ (Figure 1B and Supplemental Movie 1A), with infrequent low amplitude pulses. Addition of PD had no effect on ERKTR_C/N_, indicating that ERK activity in S1 was below the detectable limit for the reporter (Figure 1C). At low EGF concentrations (0.2-2ng/ml) we observed a range of responses, from undetectable to low amplitude pulses in ERKTR_C/N_ (Supplemental Figure 1), confirming the sensitivity of the reporter to variations in ERK activity. Addition of 20ng/ml EGF resulted in immediate and sustained activation of ERK signalling (Figure 1D, Supplemental Figure 1, and Supplemental Movie 1A). These results indicate that S1 cells depend critically on growth factor stimulation for ERK activation, with signalling dynamics sensitive to EGFR ligand concentration, similar to other non-malignant mammary epithelial cells^20^.

**Figure 1:**
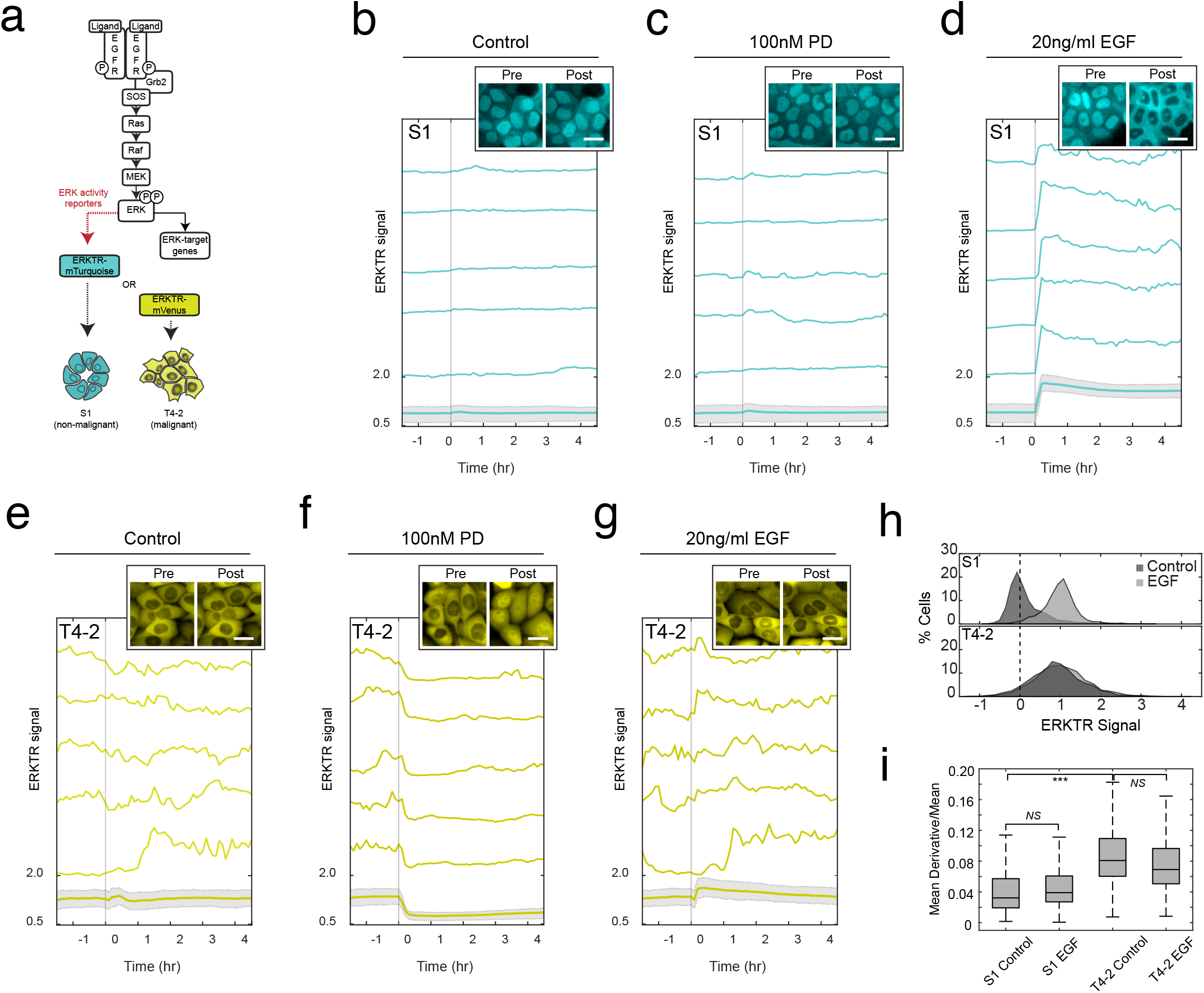
Progression to malignancy is associated with stochastic Ras/ERK signalling dynamics in a model of basal-like breast cancer. (A) Schematic of the EGFR/Ras/ERK signalling pathway in mammalian cells. ERK translocation reporters (ERKTR) carry the indicated fluorescent protein fusion to allow tracking of live-cell ERK signalling dynamics in non-malignant S1 and malignant T4-2 cells. (B) S1 (control) cells without the addition of exogenous growth factors. Bold turquoise line with gray shading represents mean ERKTR_C/N_ with the 25^th^ and 75^th^ interquartile range depicted by shading (N= 3269). Unshaded traces above represent individual cell measurements. (C) ERKTR_C/N_ following exposure of S1 cells to PD (N= 3215). (D) ERKTR_C/N_ following exposure of S1 cells to 20ng/ml EGF (N=3620). (E) T4-2 ERKTR_C/N_ signal without the addition of exogenous growth factors (N=2792). (F) Inhibition of ERKTR_C/N_ following the addition of PD to T4-2 cells (N=3075). (G) Heterogeneous induction of ERKTR_C/N_ signals in T4-2 cells treated with 20ng/ml EGF (N=2440). Dashed vertical lines indicate the addition of growth factor, inhibitor, or vehicle control at the time point indicated. For live-cell imaging examples of S1 and T4-2 control, with PD, or with EGF see Supplemental Movie 1. (H) Distributions of single cell S1 and T4-2 ERKTR_C/N_ over populations of cells with and without EGF (N= 12148). (I) Variability of ERKTR_C/N_ in control (unstimulated) versus EGF (stimulated) S1 and T4-2 cells integrated over time (N=8372). Boxes indicate 25^th^ to 75^th^ interquartile distance, and whiskers the range. NS, not significant, *P< 1e-05, ***P< 1e-150 by unpaired t-test on

In contrast to S1 cells, T4-2-ERKTR cells exhibited elevated and highly variable ERKTR signals under baseline conditions (Figure 1E and Supplemental Movie 1B). To determine their relative activation state, T4-2 cells were treated with PD, which reduced ERKTR_C/N_ to a level similar to un-stimulated S1 cells (Figure 1F and Supplemental Movie 1C). Treatment of T4-2 cells with 20ng/ml EGF provoked a maximal response similar in amplitude to EGF-stimulated S1 cells. However, the magnitude of this response relative to baseline was much smaller than in S1 due to the pre-existing high level of ERK activity in T4-2 cells (Figure 1G and Supplemental Movie 1B). Interestingly, T4-2 cells maintained stochastic signalling behavior following addition of EGF (Figure 1G), resulting in a wider distribution of ERK activities in malignant T4-2 compared to non-malignant S1 cells (Figure 1H and Figure 1I). These results indicate that progression to malignancy in HMT-3522 is associated with elevated baseline ERK signalling that is stochastic, self-sustaining, and influenced weakly by the addition of exogenous stimuli.

### Paracrine release of EGFR ligands drives ERK signalling dynamics

Paracrine/autocrine ERK activation via shedding of EGFR ligands, such as amphiregulin (Areg) and transforming growth factor-α (TGF-α) is a critical step in mammary gland development and is frequently associated with breast cancer^14, 21, 22^. TGF-α functions through autocrine mechanisms, as the high affinity of this ligand results in its immediate capture by EGFR, preventing diffusion and paracrine signalling^23^. However, Areg has a relatively low EGFR binding affinity, allowing it to diffuse freely and activate Ras/ERK signalling through paracrine mechanisms^24^. Consistent with these possibilities, treatment of T4-2 cells with EGFR inhibitors blocked Ras/ERK signalling in a dose-dependent manner and were as potent as MEK inhibitor (PD) (Figure 2A). These findings lead us to conclude that stochastic Ras/ERK signalling dynamics in T4-2 cells are dependent upon receptor-level autocrine or paracrine mechanisms, as opposed to gain-of-function mutations or other mechanisms downstream of EGFR.

**Figure 2:**
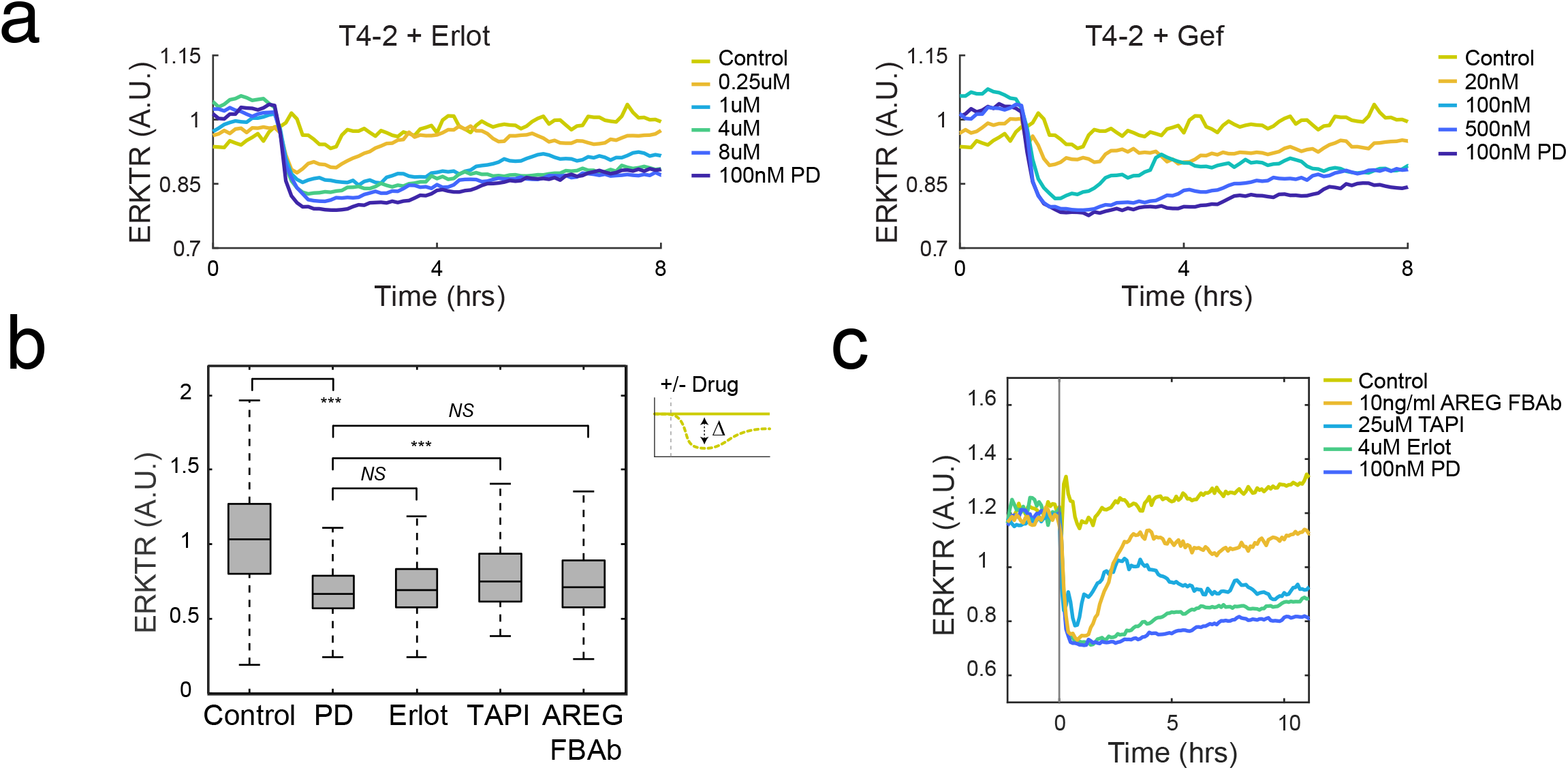
Paracrine amphiregulin release drives Ras/ERK signalling in T4-2 cells. (A) Mean ERKTR_C/N_ traces of T4-2 cells were treated with Erlotinib of Gefitinib at the indicated concentrations compared to PD. (B) Box plots depicting the maximal suppression of ERKTR signal following exposure the indicated treatments. Boxes indicate 25^th^ to 75^th^ interquartile distance, and whiskers the range (N= 14606). (C) Traces of mean ERKTR_C/N_ suppression over time for the indicated compounds (N=6116). *NS*, not significant, **P<1e-10, *** P< 1e-150 by unpaired t-test.

To determine if paracrine Areg or TGF-α account for stochastic Ras/ERK activation we tested the ability of TAPI-0, an inhibitor of the Areg and TGF-α shedding enzyme ADAM17, or Areg function blocking antibody (FBAb) to block Ras/ERK signalling. TAPI-0 inhibition produced an initial suppression of ERK activity followed by a return to dynamic signalling at a lower overall amplitude (Figure 2B and 2C p= 1.2e-17 and Supplemental Figure 2). Addition of Areg FBAb resulted in suppression of ERKTR intensity to the same extent as PD or EGFR inhibitor (Figure 2B and 2C). Similar to TAPI-0 treatment, Areg FBAb produced an initial suppression followed by a period of signalling inactivity. However, there was a gradual return to ERK dynamics comparable to control levels (Supplemental Figure 2) consistent with possible saturation of the FBAb over time resulting from continuous Areg secretion by T4-2 cells. To rule out a contribution from TGF-α, we tested high concentrations of TGF-α FBAb, which had no detectable effect on T4-2 ERKTR_C/N_ (data not shown). Based on the equivalent level of suppression achieved with EGFR inhibitors, PD, or Areg FBAb, we conclude that paracrine release of Areg is responsible for driving stochastic Ras/ERK signalling in T4-2 cells.

### Dynamic Ras/ERK signalling and target gene expression kinetics cooperatively generate expression heterogeneity

We next investigated whether the expression of ERK target genes (ETGs) involved in tumour progression is influenced by paracrine ERK dynamics. In baseline-treated T4-2 cells, we detected variable expression of the ETGs Egr1, c-Myc, Fra-1 and c-Fos, by immunofluorescence (Figure 3A) with high coefficients of variance (99%, 71%, 54%, and 62%), respectively; Fig. 3B). Co-staining of Fra-1 and Egr1 revealed discordant patterns of expression between these two proteins despite their known co-regulation by ERK signalling (Figure 3C). While this divergence could be explained by decoupling from Ras/ERK signalling in cancer, this possibility was ruled out by treatment of T4-2 cells with erlotinib, which suppressed mean Fra-1 and Egr1 expression (2.3-fold and 2.5 fold, respectively, Supplemental Figure 3A). Thus, a more likely explanation is that single-cell expression differences are driven by distinct induction and turnover parameters for these two proteins^25, 26^ and/or cell-to-cell variation in paracrine Areg-induced signalling dynamics.

**Figure 3:**
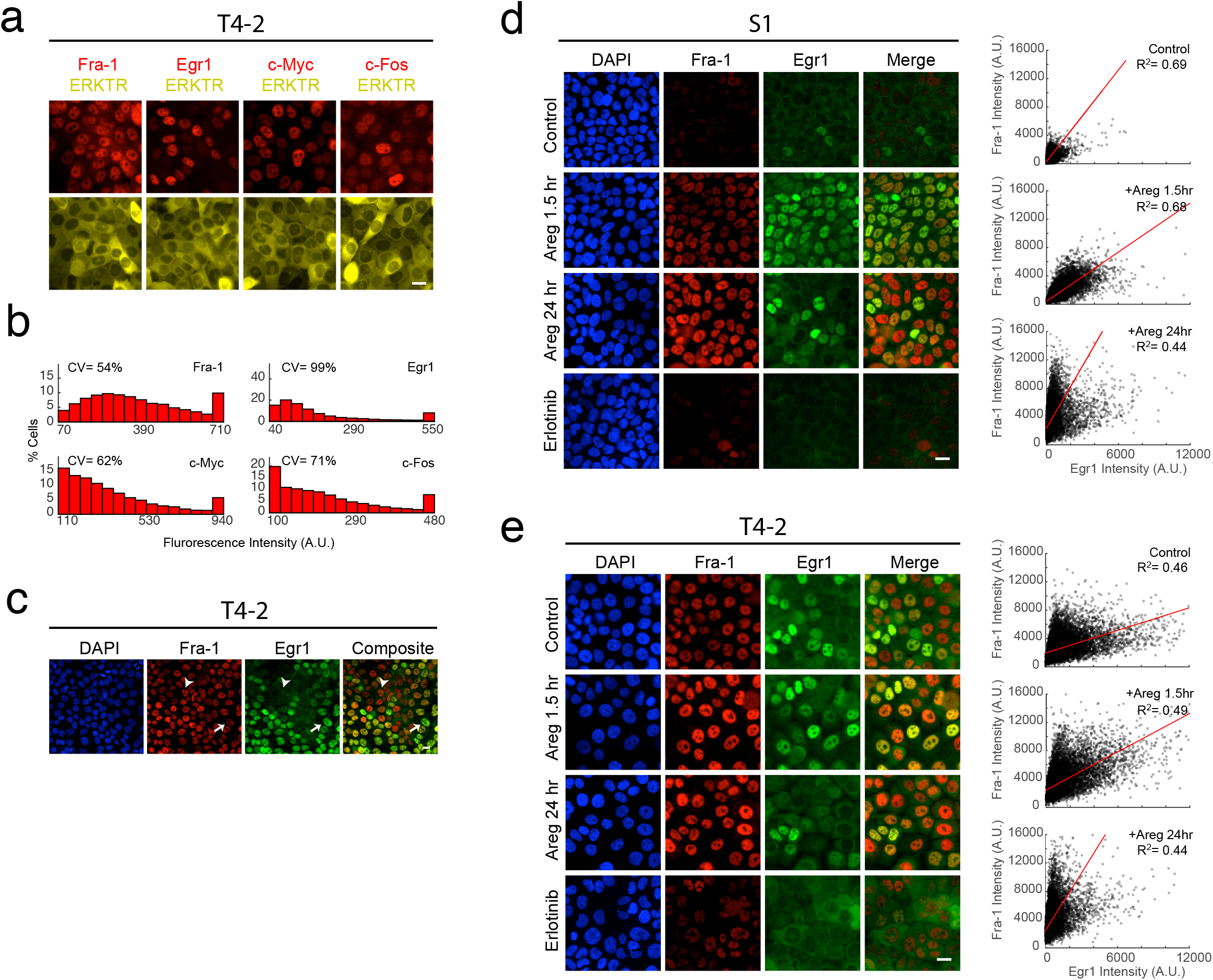
Paracrine amphiregulin drives heterogeneous ERK target gene expression. (A) Immunofluorescence of ERK target genes (ETGs) in T4-2 cells expressing the ERKTR reporter in the absence of exogenous growth factors. (B) Histograms depicting the fluorescence intensity distribution of ETG proteins under baseline conditions in T4-2 cells, N=4146 (Fra-1 and Egr1), N=9047 (c-Myc), N=7947 (c-Fos). (C) Co-staining of Fra-1 (red) and Egr1 (green) in T4-2 cells showing the co-expression profile of these proteins in signal cell. Arrowheads indicate individual cells with high Fra-1 and low Egr1 (short stem arrow) or low Fra-1 and high Egr1 (long stem arrow). (D) Time-dependent expression of Fra-1 (red) and Egr1 (green) in S1 cells following exposure to 100ng/ml Areg or 4µM erlotinib for the indicated times. Scatter plots depict single-cell Fra-1 and Egr1 signal intensities and intensity correlations (red lines) using a Theil Sen Regression analysis. Control (N=7107), 1.5 hour Areg (N=5975), 24 hour Areg (N=7172) (E) Time-dependent expression of Fra-1 (red) and Egr1 (green) in T4-2 cells following exposure to 100ng/ml Areg or 4µM erlotinib for the indicated times. Scatter plots depict single-cell Fra-1/Egr1 signal intensities and intensity correlations using a Theil Sen Regression analysis. Control (N=8947), 1.5 hour Areg (N=7030), 24 hour Areg (N=5994) Scale bars represent 20µm.

To evaluate these possibilities, we first examined the temporal response of these genes in the absence of paracrine signalling using S1 cells, which displayed low or undetectable Fra-1 and Egr1 in both control and erlotinib-treated conditions (Figure 3D). Consistent with its reported mRNA kinetics^25, 26^, Fra-1 expression gradually increased in a manner dependent upon the duration of Areg stimulus, leading to increased but heterogeneous expression levels over time (Figure 3D). Conversely, Egr1 expression peaked rapidly at 1.5 hours in all cells and became heterogeneous following prolonged stimulation with Areg (Figure 3D), consistent with its more rapid turnover^25, 26^. These findings suggest that the discordant expression of Egr1 and Fra-1 observed in T4-2 cells (Figure 3E, R^2^= 0.49) relative to unstimulated S1 cells (R^2^=0.69), results in part from the induction and turnover kinetics of individual ETGs within these asynchronously signalling cells. Similar patterns of expression were also obtained for other ETGs, including c-Fos and c-Myc (Supplemental Figure 3A) indicating that this phenomenon is shared among multiple target genes.

To test the contribution of paracrine Areg signalling to fluctuations in gene expression, we created a co-culture model using S1 and T4-2 cells carrying ERKTR sensors fused to different fluorescent proteins (Figure 4A). Since S1 cells do not secrete paracrine Areg and do not carry oncogenic mutations, by combining them with T4-2 cells we could measure directly how paracrine signalling induces ERK activity and expression heterogeneity without additional confounding factors. We co-plated cells using different S1 to T4-2 ratios and measured ERKTR_C/N_ signals (Figure 4A, Supplemental Movies 2A-G); 30:70 ratios of S1 to T4-2 produced the most reliable activation of S1 ERKTR for our assay. Under these conditions we observed an increase in S1 ERKTR_C/N_ that varied over time within each cell, compared to mono-culture conditions (Figures 4B and Supplemental Movie 2A and 2D). Addition of erlotinib reduced S1 ERKTR_C/N_, confirming that under these conditions S1 cells exhibit an elevated baseline ERK activity that is driven by paracrine signals from T4-2 cells (Figure 4C and Supplemental Movie 3B). Addition of 20ng/ml EGF increased mean ERKTR_C/N_ moderately (Figure 4D and Supplemental Movie 3A), and treatment of co-cultures with TAPI-0, Areg FBAb, or erlotinib confirmed that activation of Ras/ERK signalling in co-cultured S1 cells occurs through paracrine mechanisms (Supplemental Figure 4A).

**Figure 4:**
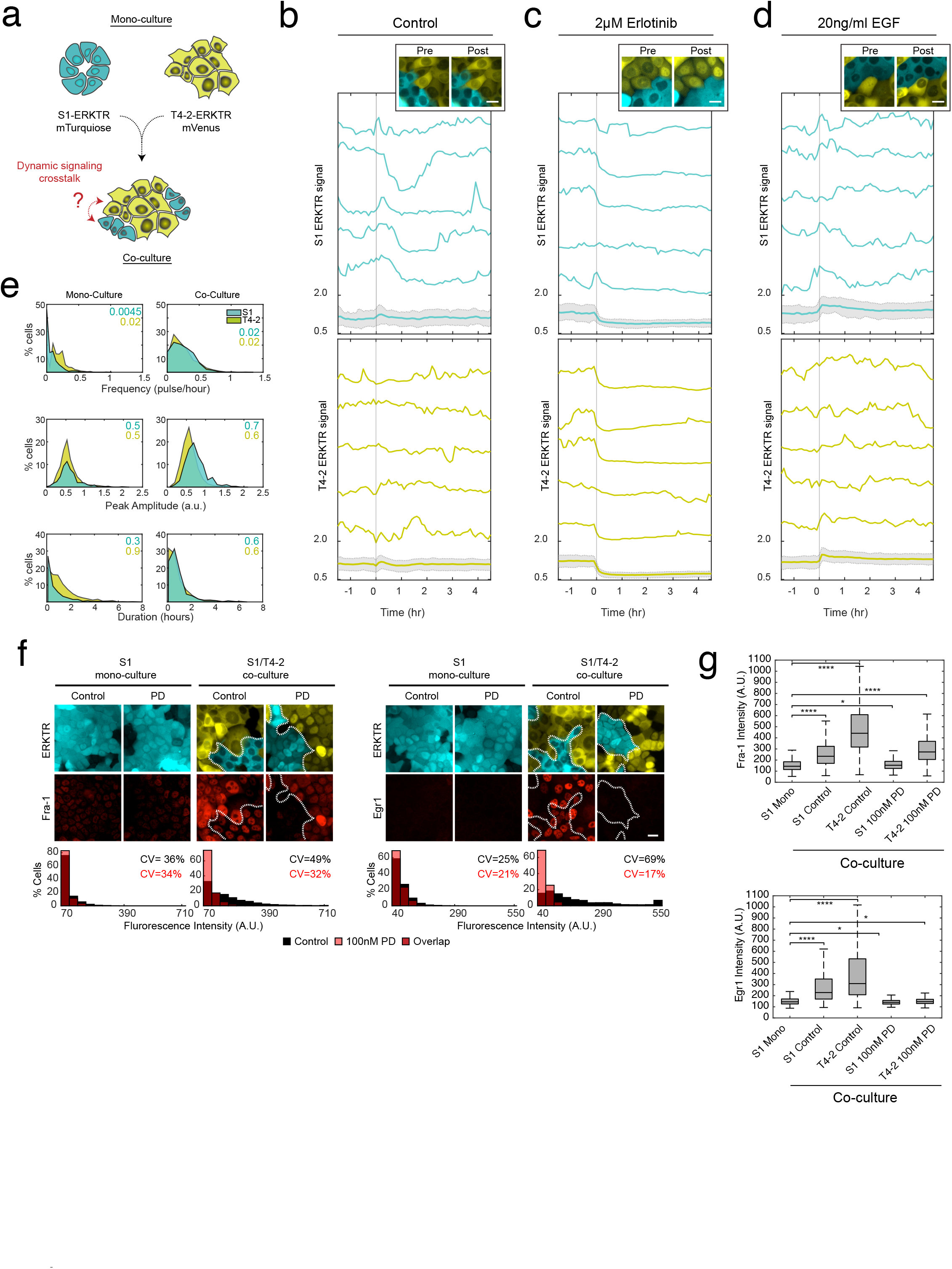
Paracrine amphiregulin signalling transfers malignant Ras/ERK signalling characteristics to proximal non-malignant cells. (A) Schematic depicting co-culture of S1 and T4-2 cells carrying ERKTR-mTurquoise or ERKTR-mVenus to visually barcode these cells for live-cell imaging, respectively. (B) S1 (turquoise lines) and T4-2 (yellow lines) ERKTR_C/N_ signals in co-culture without the addition of growth factors or inhibitors (N=1955). Bold lines with gray shading represent the mean ERKTR_C/N_ and 25^th^ and 75^th^ interquartile range. Unshaded traces represent single cell ERKTR_C/N_ measurements. (C) S1 and T4-2 ERKTR_C/N_ following treatment with 4µM erlotinib at the indicated time point (N=3539). (D) S1 and T4-2 ERKTR_C/N_ following addition of 20ng/ml EGF at the indicated time point (N=2592). (E) Distributions of single-cell pulse frequency (per hour), duration (per hour), and amplitude (arbitrary units) of ERKTR_C/N_ for S1 (turquoise) and T4-2 (yellow) under the indicated conditions (N=3497). Numbers in upper right correspond to median pulse frequency, amplitude, or duration color coded to each cell type. For live-cell imaging examples of co-cultured S1 T4-2 control, with PD, or with EGF see Supplemental Movie 3. (F) Mono-cultured S1 (turquoise) and co-cultured S1 and T4-2 (yellow) cells stained for ETG expression of Fra-1 and Egr1 (red) under control or PD conditions. Dashed lines indicate division between S1 and T4-2 cells in co-culture images. Histograms of Fra-1 and Egr1 protein expression distributions were obtained from immunofluorescence staining. Control treated data are depicted in black, PD-treated in pale red, and overlapping data in dark red (N=25169). (G) Box and whisker plots comparing mean Fra-1 and Egr1 protein levels between mono-culture S1 cells versus co-cultured S1 and T4-2 cells under control or 100nM PD treated conditions (N=30370). * P< 1e-10. ****P= 0 to machine precision.

Strikingly, co-cultured S1 cells exhibited highly variable profiles of ERK activity (Fig. 4B, D) similar to T4-2 cells under monoculture conditions (Figure 1G). Quantitatively, in co-culture conditions, S1 cells exhibited an increase in pulse frequency, amplitude, and duration similar to the stochastic pulse characteristics of T4-2 cells (Figure 4E). We evaluated the consequences of these induced ERK dynamics in S1 cells by measuring immunofluorescence of ETGs (Fra-1, Egr1, c-Myc, and c-Fos) in mono- and co-cultured cells (Figure 4F and Supplemental Figure 5A). Fra-1 and Egr1 were more broadly distributed in co-cultured S1 cells (Figure 4F; CV=49% and 69%) than in mono-cultured cells (CV=36% and 25%), approaching the variation observed in mono-cultured T4-2 cells (Figure 3A, CV=54% and 99%). Inhibition of Ras/ERK signalling reduced ETG expression and variance for these proteins (Figure 4G). Thus, in receiving cells, stochastic ERK signalling arising from paracrine presentation of EGFR ligands combines with gene-specific differences in kinetics to increase the variability of ETG expression.

### Paracrine signalling drives time-variable gene expression in surrounding non-malignant cells

A key question is whether the heterogeneity in gene expression induced by paracrine signalling reflects a population of diverse but static cell states, or time-dependent fluctuations within individual cells. To investigate this question, we utilized MCF10A cells carrying a gene fusion of mCherry at the endogenous Fra-1 locus (Fra-1::mCherry) and the ERK sensor EKAR3^20^. We first tested different ratios of MCF10A to T4-2 cells and observed a direct correlation between T4-2 density and EKAR3 signal, with the highest levels seen for 30:70 MCF10A to T4-2 ratios, similar to previous co-culture experiments (Figure 5A). Under each condition, mean Fra-1 expression remained constant, at levels corresponding with EKAR activity (Figure 5B, Supplemental Figure 5A, and Supplemental Movie 4A and 4B). While Areg treatment increased the mean Fra-1 intensity in mono-cultured MCF10A cells by 24% (Fig. 5C), relative variation remained essentially constant (CV=27% for untreated, 28% for Areg-treated, Fig. 5E). In co-culture, mean Fra-1 intensity was similar to the maximum induced by Areg (Figure 5D), but coincided with a higher degree of variation (Figure 5E, CV=35%).

**Figure 5:**
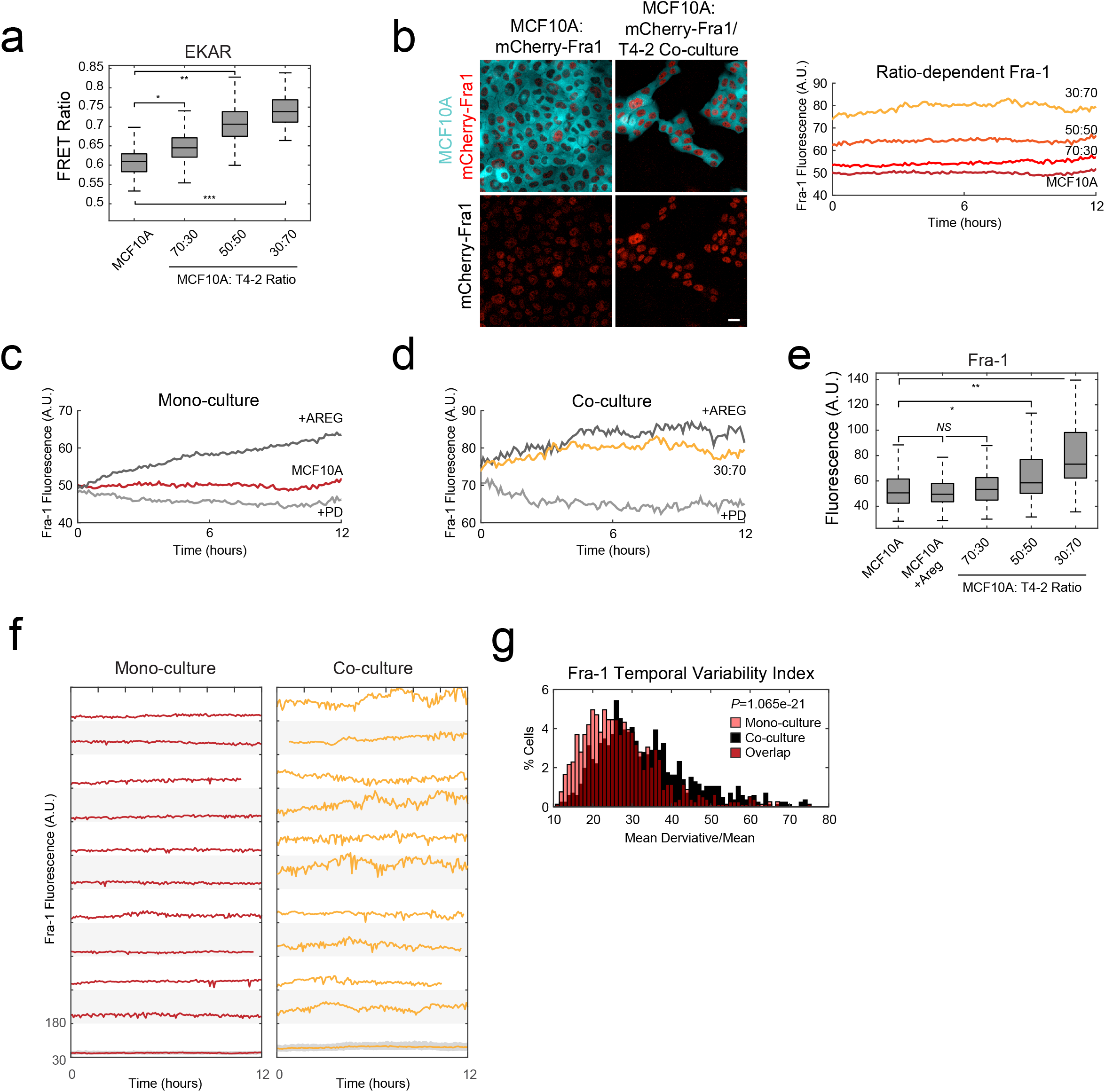
Live-cell imaging of paracrine-induced Fra-1 expression plasticity in co-culture. (A) Box plots depicting the mean FRET ratio signal in MCF10A cells carrying the ERK reporter, EKAR, in mono-culture (N=2428) and in co-culture with T4-2 at the indicated ratios (70:30 N=2526, 50:50 N=2014, 30:70 N=998). Boxes indicate the 25^th^ to 75^th^ interquartile distance and whiskers the range. (B) Still frames of MCF10A EKAR Fra1::mCherry cells in mono-culture and co-culture captured by live-cell imaging. Upper panels depict EKAR expression (mTurquoise2, cyan) and tagged endogenous Fra-1 (mCherry, red); T4-2 cells are not shown in co-culture images. Lower panels show Fra-1::mCherry expression only (red). Scale bar represents 20µm. The graph depicts mean Fra-1 intensity over time for each ratio of T4-2 co-culture (MCF10A, 70:30, 50:50, and 30:70 MCF10A to T4-2 ratios) as labeled on the graph. Culture media did not contain exogenous growth factors. (C) Fra-1::mCherry expression in mono-cultured, unstimulated MCF10A cells (dark red, re-plotted from panel B), with exogenous Areg at 100ng/ml (dark gray), and 100nM PD (light gray). (D) Fra-1::mCherry expression in MCF10A cells co-cultured at a 30:70 ratio of MCF10A to T4-2 cells (orange, re-plotted from panel B), in comparison to MCF10A cells at the same co-culture ratio treated with 100 ng/ml exogenous Areg (dark gray), or 100nM PD (light gray). See Supplemental Movie 4 for live-cell examples. For panels B through D, mono-cultured MCF10A (N=786), +Areg (N=1560), +PD (N=544), co-cultured 70:30 (N=945), 50:50 (N=1028), 30:70 (N=865), +PD (N=622). (E) Box plots depicting the mean Fra-1 expression levels under the indicated conditions. Boxes indicate the 25^th^ to 75^th^ interquartile distance and whiskers the range. MCF-10A (N= 157), 70:30 (N= 182), 50:50 (N= 160), 30:70 (N= 66). (F) Fra-1::mCherry fluorescence over time (12 hours) in individual mono-cultured (red) or co-cultured (30:70 ratio) MCF10A cells (orange). The bottom trace represents mean Fra-1 signal with the interquartile range shaded, the traces above depict representative single cell traces of Fra-1 expression over time (Mono-culture N=786, Co-culture N=865). (G) Distribution of temporal variability index in individual cells, defined as the absolute value of the derivative Fra-1 signal/mean Fra-1 signal for each cell.

At the single-cell level, temporal fluctuations in Fra-1 expression were evident under co-culture conditions, in comparison to mono-cultured cells (Figure 5F). To rule out the possibility that this heterogeneity is simply a function of high frequency noise, we filtered our data (see Methods) to focus on gradual changes in Fra-1 over time (>30mins). We then calculated the mean derivative, normalized by the mean Fra-1 intensity on a single-cell basis, to obtain a measure of the variance of Fra-1 expression over time (Temporal Variability Index) in mono-culture versus co-culture. By this measure, variability in Fra-1 increased by 20% in co-culture relative to mono-culture (Figure 5G, p=1.065e-21). Thus, the observed heterogeneity in Fra-1 expression reflects not only static variation across a population but time-varying effects within single cells. Notably, because Fra-1’s mRNA makes it resistant to temporal variation in ERK activity^20^, these conditions would be expected to have even larger temporal effects on other ETGs.

### Paracrine signalling crosstalk diversifies global gene expression

Because many ETGs, including Fra-1, c-Myc, and Egr1, are themselves transcription factors, fluctuations in their protein levels could impact an even larger set of genes, resulting in global variation of transcriptional profiles over time. To capture this possible diversity, we performed single cell mRNA sequencing (scRNAseq) of S1 (expressing ERKTR:mCherry) and T4-2 (ERKTR:mVenus) cells in mono and co-culture using the DropSeq method^27^ (Figure 6A). For co-cultured cells, sequencing reads of mCherry and mVenus tags were used to classify cells without the need for physical sorting (see materials and methods). We performed cluster analysis of the cells (see materials and methods) from mono-cultured S1 controls (S1), S1 treated with 100ng/ml Areg (S1a), and T4-2 (T4) and found that cells broadly sort according to class and treatment (Figure 6B), validating our ability to resolve each cell class with this method. Similarly co-cultured S1/T4-2 (labeled S1co and T4co, respectively) cells segregated into distinct clusters, even without filtering by expression of mCherry or mVenus (Supplemental Figure 7A and Figure 6B (only S1co shown for clarity)). At the selected resolution, we recovered eight clusters of cells with similar transcriptional signatures (Figure 6C and Supplemental Data 1). We performed differential expression analysis to determine gene sets enriched in each cluster. T4-2 cells comprise 3 clusters, enriched for genes involved in replication and cell division (cluster 3) and a diverse array of genes including AREG, reactive oxygen enzyme SOD2, and TNF-related Apoptosis Inducing Ligand TRAIL/TNFSF10 (clusters 0 and 7) (Figure 6C and 6D). S1 cells treated with Areg (S1a) displayed an increase in the expression of cell-cycle associated genes, consistent with this ligand acting as a mitogen (cluster 5), whereas the bulk population of cells (cluster 1) showed broad changes in gene expression with an increase in ERK regulated genes (e.g. c-Myc) compared to unstimulated controls. Interestingly, S1co cells formed a separate cluster (6) with an expression profile that partially overlapped with monocultured S1 and T4-2 (Figure 6C and 6D), suggesting that they enter a hybrid transcriptional state under these conditions.

**Figure 6:**
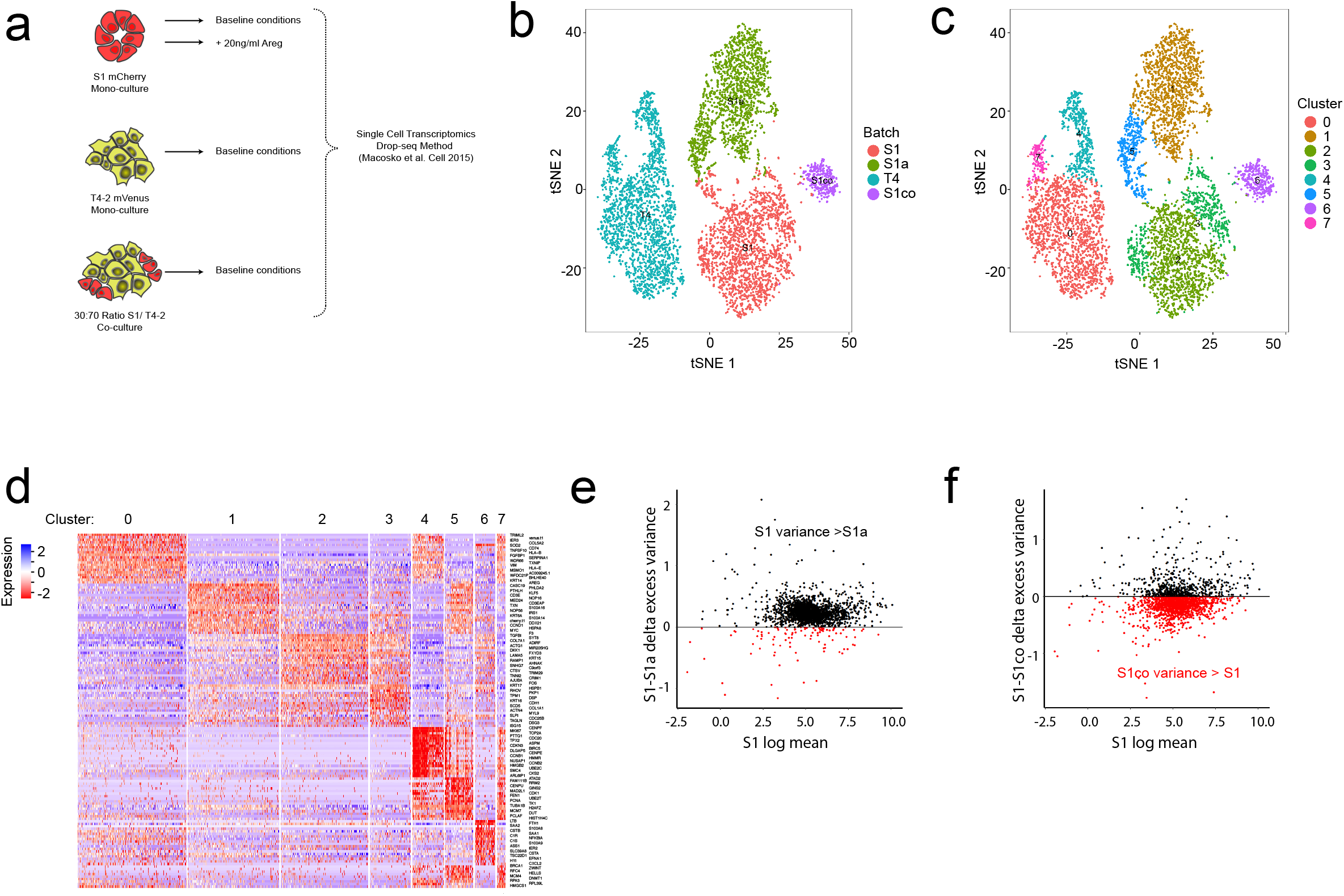
Induction of global gene expression variance by cellular cross-talk. (A) Schematic depicting cell culture conditions for S1, T4-2, and S1/T4-2 co-cultures for single-cell RNA sequencing. Red: S1 cells expressing ERKTR-mCherry; yellow: T4-2 cells expressing ERKTR-mVenus. (B) tSNE plot of S1 control (orange, N=2252), S1 treated with 20ng/ml amphiregulin (S1a, green, N=1920), T4-2 control (blue, N=2493), or S1 (purple, N=342) co-cultured with T4-2 cells (not shown, N=2493), clustered based on single cell transcript profiles by cell type. (C) tSNE plot with 8 clusters representing distinct transcriptional profiles. (D) Heatmap of top 20 biomarkers over-represented in cells of a cluster. Numbers correspond with the clusters indicated in (C). (E) Difference in excess gene expression variance between S1 control cells (S1) and S1 treated with amphiregulin (S1a) plotted against the log mean S1 variance. Dots represent a single gene from ~1900 genes sampled. Red dots represent genes with an increased excess variance in S1a compared to S1. Black dots represent genes displaying a decreased excess variance in S1a compared to S1 (P = 6.2×10^-271^). (F) As in (E), the difference between excess variance is plotted for S1 control (S1) and S1 co-culture (S1co) against the log mean S1 variance. Red dots depict genes with an increased excess variance in S1co compared to S1. Black dots represent genes with a decreased variance in S1co compared to S1 (P = 1.7×10^-61^).

Next, we aimed to determine if gene expression exhibits a globally heterogeneous (i.e. variable from cell-to-cell) response to these conditions similar to our local observations for ETG proteins alone (see Figure 3). We calculated the excess variance in gene expression between different cells and treatments using log normalized data, where excess variance is measured as the variability of gene expression beyond a fitted mean-variance relationship. We predicted that brief treatment of S1 cells with exogenous Areg (t <2 hours) would reduce excess variance by saturating ERK-dependent expression responses^28^ (see Figure 3E time course), whereas S1co cells, which were exposed to more stochastic levels of Areg, would display an increase in variance. Indeed we found that treatment of cells with 100ng/ml Areg, led to a reduction in gene expression variance compared to S1 control cells (Figure 6E, P = 6.2×10^-271^). By contrast, S1 cells when co-cultured with T4-2 cells display an increase in excess variance (Figure 6F, P = 1.7×10^-61^) compared with S1 in monoculture, indicating that these conditions favor the induction of heterogeneous expression profiles from cell to cell. Additionally, we found that in co-culture, S1 cells also had a substantial effect on the T4-2 gene expression profile. We found that T4-2co cells formed clusters distinct from mono-cultured T4-2 cells (Supplemental Figure 7A and 7B) and that the presence of S1 cells significantly increased the gene expression variance of T4-2 cells (Supplemental Figure 7C). These findings indicate that bidirectional interactions of different cell types favor the diversification of gene expression and implicate signalling cross-talk as a driver of this process.

## Discussion

By exploring diverse cellular phenotypes, tumour cells gain a selective survival advantage in adverse physiological environments^29^. Genetic mutation and epigenetic modulation of gene expression are important drivers of tumour cell diversification, but are constrained to operate on the timescales of weeks or longer. Immediate cell survival in response to stress conditions may require a more rapid means of phenotypic diversification. We demonstrate here that continual variation in growth factor signalling throughout the TME, transduced through Ras/ERK signalling, is sufficient to provoke time-dependent fluctuation of ERK target genes, including Fra-1, a transcription factor linked to metastasis and drug resistance in multiple cancer types (Figure 7). While this system is highly simplified relative to actual tumours, these data establish the principle that temporal plasticity in the expression of key tumour genes can be directly driven on the time scale of hours by factors known to exist within the TME.

**Figure 7:**
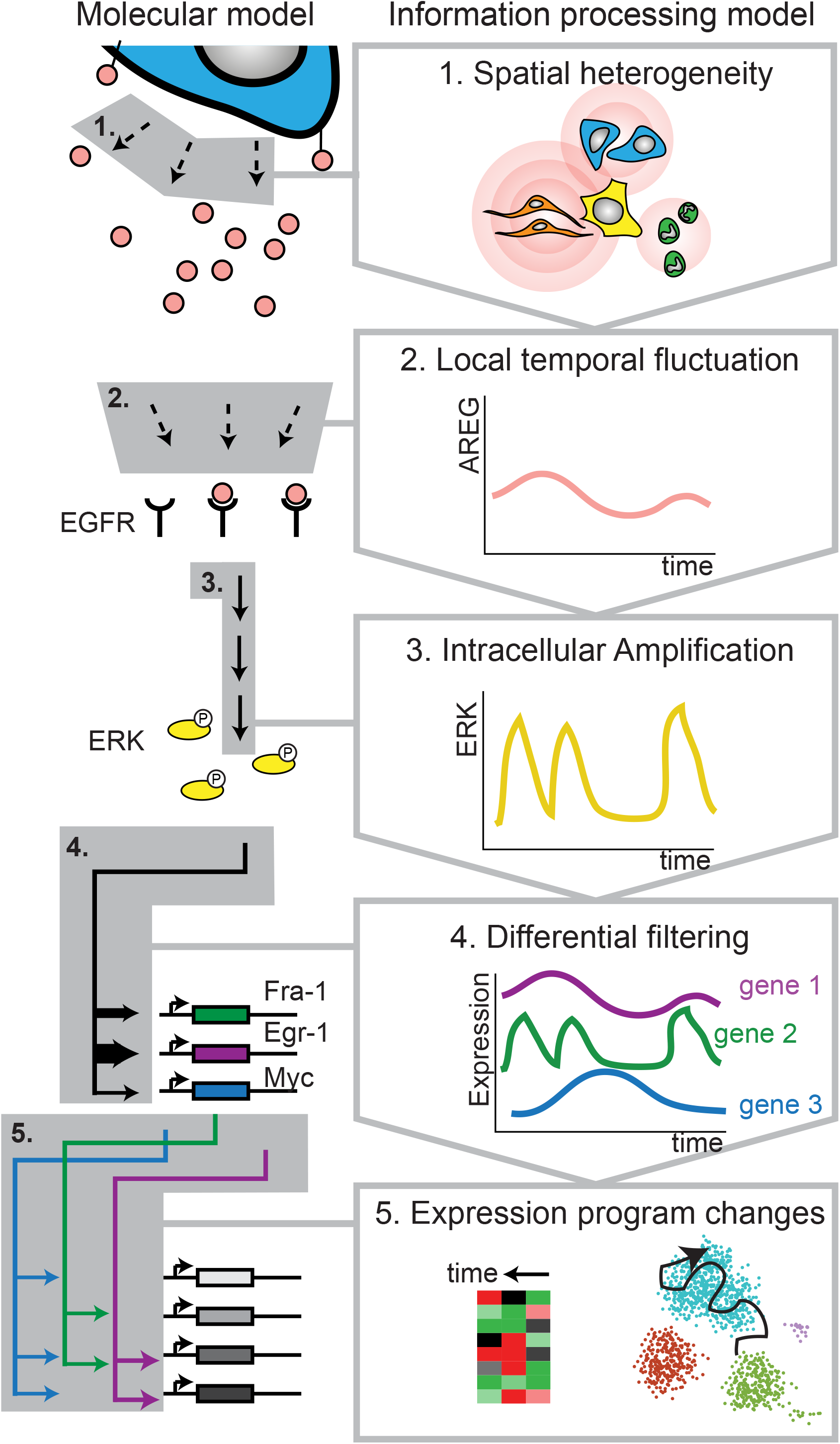
A plasticity generator model. Molecular processes are shown at left, and the information processing function performed by each biochemical step is shown at right. 1. Physical arrangements of different cell types vary throughout a tumour, resulting in non-uniform secretion and micron-scale concentration gradients of EGFR ligands such as Areg. 2. The convergence of multiple Areg gradients at the surface of a receiving cell results in fluctuations over time in the locally available Areg concentration. 3. The EGFR/Ras/ERK cascade acts as a nonlinear amplifier that converts extracellular fluctuations in Areg into high-amplitude pulses of ERK activity. 4. The multiple target genes of ERK respond differentially to the sequence of activity pulses, leading to non-synchronous fluctuations in their protein expression. 5. The activities of fluctuating ETG proteins as transcription factors results in fluctuation in the expression of other genes (left), the net result of which is a more processive exploration of gene expression space (right). This traversal of gene expression states provides cells with phenotypic plasticity.

The mixture of malignant (T4-2) and non-malignant (S1 or MCF10A) cells within our system represents an approximation of the genetic diversity within a tumour, where clones containing different mutational burdens would interact. Our experiments provide insight into the behavior of both types of cells. The proficiency of T4-2 cells to self-stimulate through paracrine mechanisms is consistent with previous work^19, 30^, and we find that this mode of activation of Ras/ERK signalling induces both a high amplitude of ERK activity as well as time-dependent variation that is a likely source of the heterogeneity we observe at the level of gene expression. Conversely, co-cultured non-malignant cells acquire characteristics of Ras/ERK activity and ETG expression similar to malignant cells, even in the absence of activating mutations in the Ras/ERK pathway. Within our experimental system, the low level of constitutive ERK signalling and high capacity for stimulation of these cells makes them ideal “receivers” to quantify the paracrine stimuli emanating from the nearby malignant cells. An intriguing but unresolved question is whether, within the context of a real tumour, such “receiver” cells represent a population that is important for tumour progression, or if such cells (e.g. non-malignant cells) can be reprogrammed by these signals to take on a malignant phenotype. In principle, such cells, supported by the signals from more highly mutated cells nearby, could represent a reservoir of uncommitted possibilities with higher adaptive potential than heavily mutated cells that cannot tolerate further genetic alterations. Such flexibility could play a role in tumour cell survival when new stresses, such as cytotoxic chemotherapeutics, are encountered.

The general process we demonstrate, by which spatial heterogeneity in paracrine signalling drives temporal variability in gene expression, may also apply in other signalling pathways, such as IL-6^31, 32^ or TGF-β^33, 34^ signalling. However, we note that the kinetic parameters of the amphiregulin-EGFR-Ras-ERK pathway are especially well suited for generating global diversity in gene expression (Figure 7). The amplification characteristics of the EGFR-Ras-ERK pathway are capable of driving high-intensity ERK activity bursts in response to even small amounts of ligand, and the attenuation of these pulses by negative feedback is rapid, resulting in ERK output that magnifies external changes in amphiregulin concentration. Subsequently, the genes responding to ERK activity do so with a wide range of kinetic properties^26^, continuously translating the temporal dynamics of the pathway into different gene expression profiles. It is intriguing to speculate that the prevalence and potency of the Ras/ERK pathway in tumourigenesis may be a consequence not only of its target functions (as other, less prevalent pathways, also stimulate cell proliferation and migration), but also of its ability to act as a generator of cellular diversity.

Additional sources of variability, including other cytokines, extracellular matrix, and hypoxia, are certainly present within the TME, and genes other than Fra-1 contribute to tumour cell survival. Although we have not recapitulated this complexity in its entirety, our simplified system has made it possible to quantitatively characterize and model the generation of gene expression variability *in vitro*. This experimental system could be used to evaluate candidate inhibitors for their ability to reduce tumour cell plasticity, a property that is likely closely related to their efficacy^35^, but for which no direct measurement has yet been available.

## Methods

### Cell culture

HMT-3522 cell lines were maintained in DMEM/F12 (Gibco) supplemented with prolactin, insulin, sodium selenite, hydrocortisone, β-estrogen, transferrin, and EGF (S1 only), as previously described^14^. MCF10A-5E cell lines were maintained in DMEM/F12 media supplemented with 5% horse serum, insulin, cholera toxin, hydrocortisone, and EGF. All cell lines were maintained in 5% C02 at 37°C.

### Reporter line construction

Reporter cell lines were created using lentiviral transduction from plasmid DNA containing the ERK Translocation Reporter (ERKTR)^15^ fused to fluorescent proteins as indicated below. T4-2 reporter lines were constructed with ERKTR-mVenus. S1 reporter lines were constructed using ERKTR-mTurquoise or ERKTR-mCherry. Reporter expressing cells were selected with puromycin followed by sorting by flow cytometry. Flow sorting was conducted using a wide gating strategy to maximally preserve the inherent heterogeneity of S1 and T4-2 cell populations. MCF10A-EKAR-Fra-1::mCherry cell lines were generated previously using the EKAR3 reporter inserted into MCF10A cells carrying mCherry fused to the C-terminus of endogenous *FOSL1* (Fra-1) locus^20^.

### Live-cell microscopy and co-culture conditions

Live-cell microscopy was conducted on multi-well plates with #1.5 glass bottoms (In Vitro Scientific, Mountain View, CA) that were coated with laminin-111 (Invitrogen, Carlsbad, CA). 50ug/ml Laminin-111 was deposited onto glass wells overnight in 20mM sodium acetate buffer containing 1mM CaCl_2_ to generate a fractal laminin-coated surface^36^. Immediately prior to plating, the wells were washed once with PBS to remove excess buffer and laminin. For mono-culture experiments, cells were plated at a density of 9000 cells/well for both S1 and T4-2 cells. For co-culture experiments, total cell density was kept constant and the ratio of S1:T4-2 cells adjusted for a particular experiment (e.g. 30:70 S1 to T4-2 ratio corresponds to a combination of 2700 and 6300 cells/well, respectively). Plates were then incubated for 24 hours in complete media. After 24 hours, plates were washed twice in media containing no additives then placed in custom imaging media (DMEM/F12 without phenol red, riboflavin, and folic acid) containing hydrocortisone, β-estrogen, transferrin, and Hoecsht stain, then incubated overnight prior to live-cell microscopy. Following preparation, plates were imaged on a Nikon Ti-E inverted microscope fitted with an environmental chamber. A single stage position was chosen within each well of the plate and time lapse images were captured every 6 minutes under the indicated conditions with 20X 0.75 NA objective and Andor Zyla 5.5 scMOS camera. Automated imaging was performed using NIS-Elements AR software.

### Immunofluorescence microscopy

For fixed staining experiments cells were plated exactly as describe for live cell imaging experiments. Following treatment and incubation under the indicated conditions cells with fixed in 4% paraformaldehyde in phosphate buffered saline for 20 minutes. Wells were then blocked with buffer containing 0.1% Triton and 4% bovine serum albumin. Primary antibodies, Mouse anti-Fra-1 (Santa Cruz, sc-28310), Rabbit anti-Egr1 (Cell Signalling Technologies, #4153), Rabbit anti-c-Fos Cell Signalling Technologies, #2250), and Rabbit-anti-c-Myc (Cell Signalling Technologies, #5605) were used at 1:400 dilution. For visualization, secondary Goat anti-Mouse Alexa Fluor 647 or Goat anti-Rabbit Alexa Fluor 546 were used at a 1:200 dilution.

### Image, data processing, and statistics and normalization

For all experiments >500 cells were analyzed per condition unless otherwise indicated. Each imaging dataset presented is representative of at least two independent replicate experiments. Image, data processing, and statistics were performed using custom MATLAB software as previously described ^37, 38^. Software will be made available upon request. Linear regression was performed using a Theil-Sen estimator as previously described (ref). Temporal Variability Index (*TVI*) was calculated by taking the absolute value of the mean differential of filtered Fra-1 intensity (*Fra1_f_* over time divided by the mean.

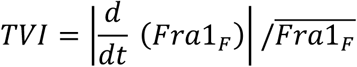

Fra-1 filtering was performed using a custom MATLAB script employing an infinite input response and Butterworth filter design set to lowpass filtering to remove high frequency noise with a periodicity < 30 minutes. All t-tests were calculated using an unpaired approach, significance was considered *P* <0.05.

### Cell lines used for sequencing and treatments

To conduct single cell RNA sequencing experiments the following cells were grown under standard culture conditions for 4 days then dissociated with 0.25% trypsin/EDTA and rinsed into phosphate buffered saline immediately before DropSeq. S1 and T4-2 mono-culture cells were grown as described previously with the media changed daily. S1/T4-2 cells were grown under the same conditions except that no exogenous growth factors were added. S1a cells received new media daily containing 100ng/ml amphiregulin. Species mixing experiments were performed using mouse embryonic fibroblasts (MEFs).

### DropSeq

Droplets were generated using a DSQ 3×9 array microfluidic part (Nanoshift, Emeryville, CA, USA) using a Drop-seq setup according to Macosko et al. 2015 (Online-Dropseq-Protocol-v.-3.1-Dec-2015, http://mccarrolllab.com/dropseq/). Droplet size was determined using fluorescent beads (P=S/2% 10µM Dragon Green, Bangs Laboratories) as described (Measuring-Droplet-Volume-in-Home-Made-Devices, http://mccarrolllab.com/dropseq/). Barcoded beads (Barcoded Beads SeqB; ChemGenes Corp., Wilmington, MA, USA) and cells were loaded at concentrations specified in Supplemental Table 1. Prior to the collection, cell syringes and tubing were blocked using PBS + 0.1% BSA. A magnetic mixing disc was inserted into the cell syringe to allow for manual cell mixing during the run and the cell pump was used in a vertical position. Droplets were collected in 50mL Falcon tubes and the target volume of aqueous flow varied in between 1-1.2mL of the cell suspension. Droplets were broken immediately after collection and reverse transcription, exonuclease-treatment and further processing was conducted as described previously ^27^. For each library, three test PCRs (50µl) each containing a bead equivalent of 100 STAMPs were conducted to determine the optimal cycle number for library amplification. 35µl of each test PCR were purified using Agencourt AMPure XP beads (Beckman Coulter, A63881, 21µl beads (0.6X) and 7µl of H_2_O for elution) and the DNA concentrations were determined using a Qubit 4 Fluorometer. A concentration in between 400-1000pg/µl was taken as optimal. A variable number of PCR reactions was conducted to amplify the available 1st strand cDNA also with 100 STAMP bead equivalents per reaction with the optimal cycle number (50 µl volume; 4 + 8-11 cycles, Supplemental Table 1). 12 µl fractions of each PCR reaction were pooled, then double-purified with 0.6X volumes of Agencourt AMPure XP beads (Beckman Coulter, A63881) and eluted in H_2_O using 1/3rd of the bead volume. 1µl of the amplified cDNA libraries were quantified Qubit 4 Fluorometer and library size distribution verified on a BioAnalyzer High Sensitivity Chip (Agilent). 600 pg cDNA of the library was fragmented and amplified (12 cycles) using the the Nextera XT (Illumina) sample preparation kit. The libraries were double-purified with 0.6x volumes of AMPure XP Beads and quantified.

### Sequencing strategy

Nextera libraries were sequenced on Illumina Nextseq 500 sequencers using the NextSeq High Output v2 kit (75 cycles), using a custom primer and a custom paired end sequencing strategy, R1 20bp, index 8bp, R2 remaining bp^27^. The five Nextera libraries were pooled and a total of 9.9k anticipated STAMPs were loaded on the flow cell with the following library portions; S1/MEF: 5%, T4: 17.1%, T4/S1: 30.3%, S1: 17.2%, S1 amphiregulin: 13.1%. The pool was sequenced on two NextSeq 500 runs. Raw sequencing data have been submitted to the GEO repository and available under GEO accession GSE118312.

### Reference for read alignment

For the species mixing experiment we used a mixed reference (human+mouse) available at GEO accession GSE63269^27^. For mapping of T4 and S1 reads we used the human genome primary assembly (GRCh38) available at https://www.ensembl.org/ and release 92 gene models. Sequences for transgenic markers mCherry and mVenus were added to the reference.

### Drop-seq pipeline and generation of the gene expression matrix

All Drop-seq data was processed using Drop-seq Tools v1.12 as described previously (http://mccarrolllab.com/wp-content/uploads/2016/03/DropseqAlignmentCookbookv1.2Jan2016.pdf)^27^.

Bowtie2 (v2.2.6)^39^ was used for read alignment with the settings —phred33 —very-sensitive -N 1.

### Species mixing experiment

To determine whether single cells were generated, we mixed S1 cells and MEF (mouse) cells, then mapped the resultant data against a hybrid reference containing both human and mouse genes. Cells were loaded at approximately 100 cells/uL (50cells/µl final), which would be expected to give ~2.5% doublet rate, similar to the 1.6.% doublet rate observed in the experiment (Supplemental Figure 8A SpeciesMix). This library was sequenced very shallow since its sole purpose was doublet assessment.

### Cell QC and clustering

Cells with more than 500/2k and less than 6k/30k genes/UMIs were considered in the analyses (Supplemental Figure 8B DropseqStats). Cells with more than 10% of mitochondrial expression were filtered out and excluded from downstream analyses. 10,212 cells remained after filtering with medians of 2,653 genes and 6,283 UMIs. Analyses were conducted using R package Seurat (v2.2.1)^40^. We provide two R markdown files with the complete analysis code. The initial clustering is described in the supplementary files (T4S1_init_clust.Rmd), subsequent analyses are provided in supplementary file X (T4S1_analyses.Rmd). We also provide knitted html versions.

### Excess variance calculation and delta-excess variance

UMI counts for each cell were converted to transcripts per million (TPM) by scaling all cell-wise counts to 1,000,000. For each cell population *i* and each gene *g*, under the assumption of a negative binomial distribution, we computed its mean expression *μ_i,g_* and variance *v_i,g_*, then calculated gene-wise dispersion parameters *α_i,g_* from the expression 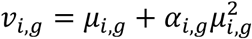. We observed a linear trend between log *μ_i,g_* and log *α_i,g_* for all cell populations, indicating as the mean expression of a gene increased, the overdispersion tends to decrease. Therefore, to measure change in dispersion for each gene between two populations (when the mean may also change), we instead compare “excess variance”. For every pair of populations *i* and *j* being compared, we fit an “excess variance” term *E_i,g_* = (*μ_i,g_* – *μ̂_i,g_*), where the fitted overdispersion estimate *μ̂_i,g_* is generated by performing Local Polynomial Regression Fitting (loess) using R version 3.4.4 with span=1, simultaneously regressing log *α_i,g_* on log *μ_i,g_* and log *α_j,g_* on log *μ_j,g_* over all genes *g*. Statistical significance was calculated using a paired, two-sided Wilcoxon signed rank test between all *E_i,g_* and *E_j,g_*. See Supplemental Data 2 for a complete list of genes and relative excess variance.

### Code Availability

All custom MATLAB and R code will be made available upon request.

## Supplemental Figure Text

**Supplemental Figure 1:**
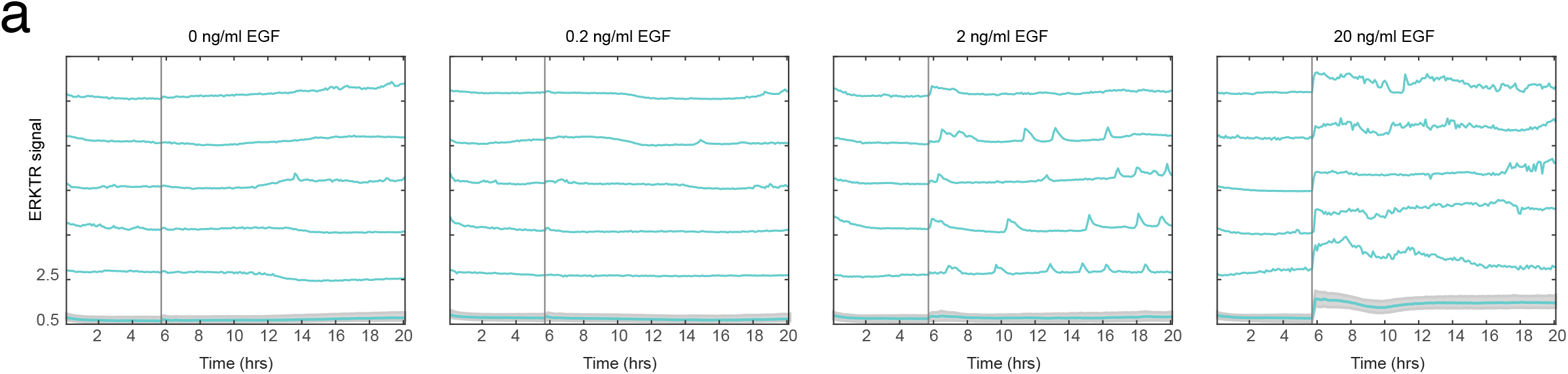
Dose-dependent EGF response in S1 cells by single cell analysis. (A) Single cell traces of S1 cells treated with the indicated concentrations of EGF for ~20 hours. Vertical line indicates the addition of EGF. Bold lines with gray shading represent the mean ERKTR response and the 25^th^ and 75^th^ interquartile range, respectively. Unshaded turquoise lines represent the ERKTR signal obtained from individual cells.

**Supplemental Figure 2:**
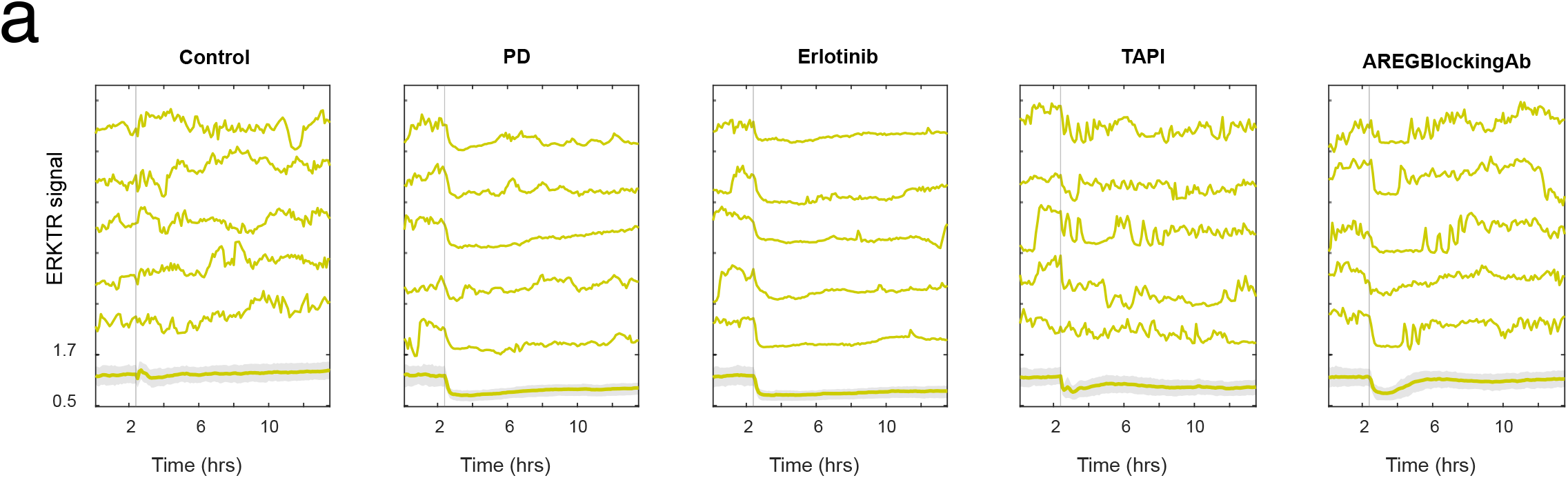
Compound-specific Areg inhibition kinetics. (A) Mean and single cell traces of T4-2 cells treated with 100nM PD, 4µM erlotinib, 25µM TAPI, or 10ng/ml AREG function blocking antibody compared to control. Bold lines with gray shading represent the mean ERKTR response and the 25^th^ and 75^th^ interquartile range, respectively. Unshaded yellow lines represent the ERKTR signal obtained from single cells.

**Supplemental Figure 3:**
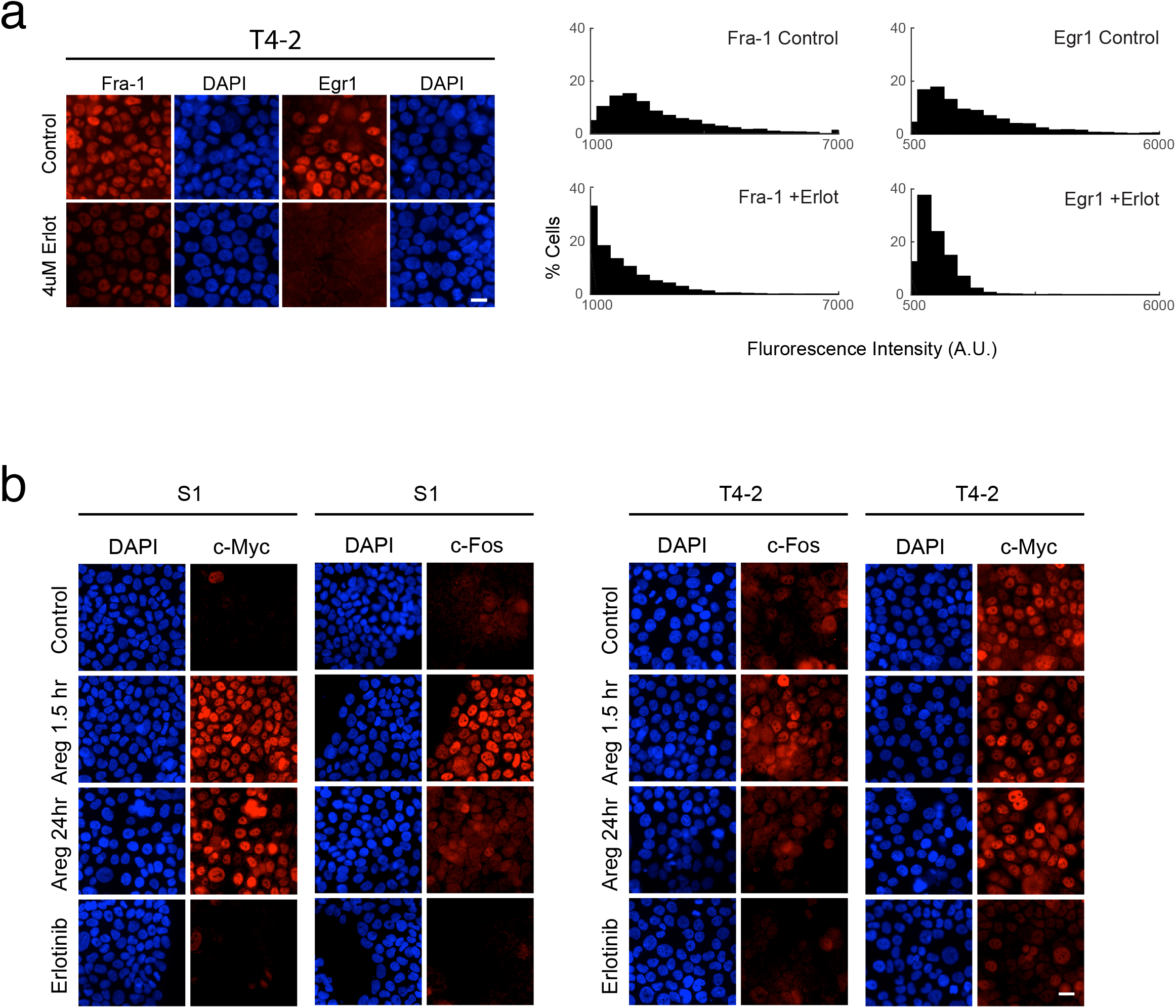
Temporal expression characteristics of c-Fos and c-Myc in response to exogenous growth factor or EGFR inhibition: (A) Comparison of Fra-1 and Egr1 levels in T4-2 cells with or without 4µM erlotinib. (B) Immunofluorescence images of S1 and T4-2 cells treated with 100ng/ml Areg or 4µM erlotinib for the indicated time periods. DAPI is shown in blue, c-Fos and c-Myc are shown in red. Scale bar represents 20µm.

**Supplemental Figure 4:**
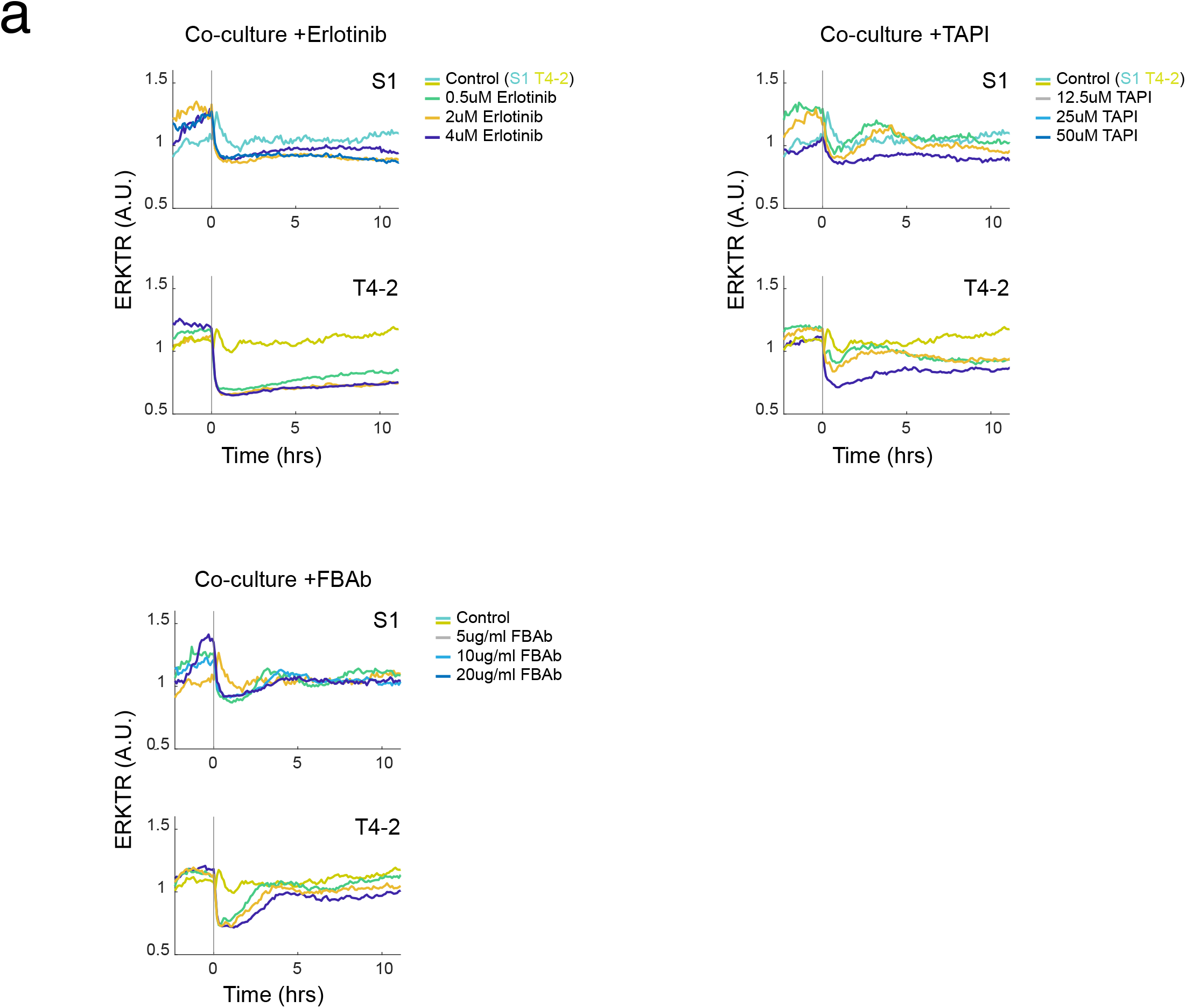
Dose-dependent inhibition of paracrine signalling in S1/T4-2 co-cultures. (A) Mean ERKTR signal in co-cultured S1 and T4-2 treated with 0.5, 2, or 4µM erlotinib over indicted time course. (B) Mean ERKTR signal in co-culture cells treated with 12.5, 25, or 50µM TAPI. (C) Mean ERKTR signalling in co-cultured cells treated with 5, 10, or 20ng/ml Areg function blocking antibody. Vertical line depicts the time of erlotinib, TAPI, or Areg function blocking antibody addition.

**Supplemental Figure 5:**
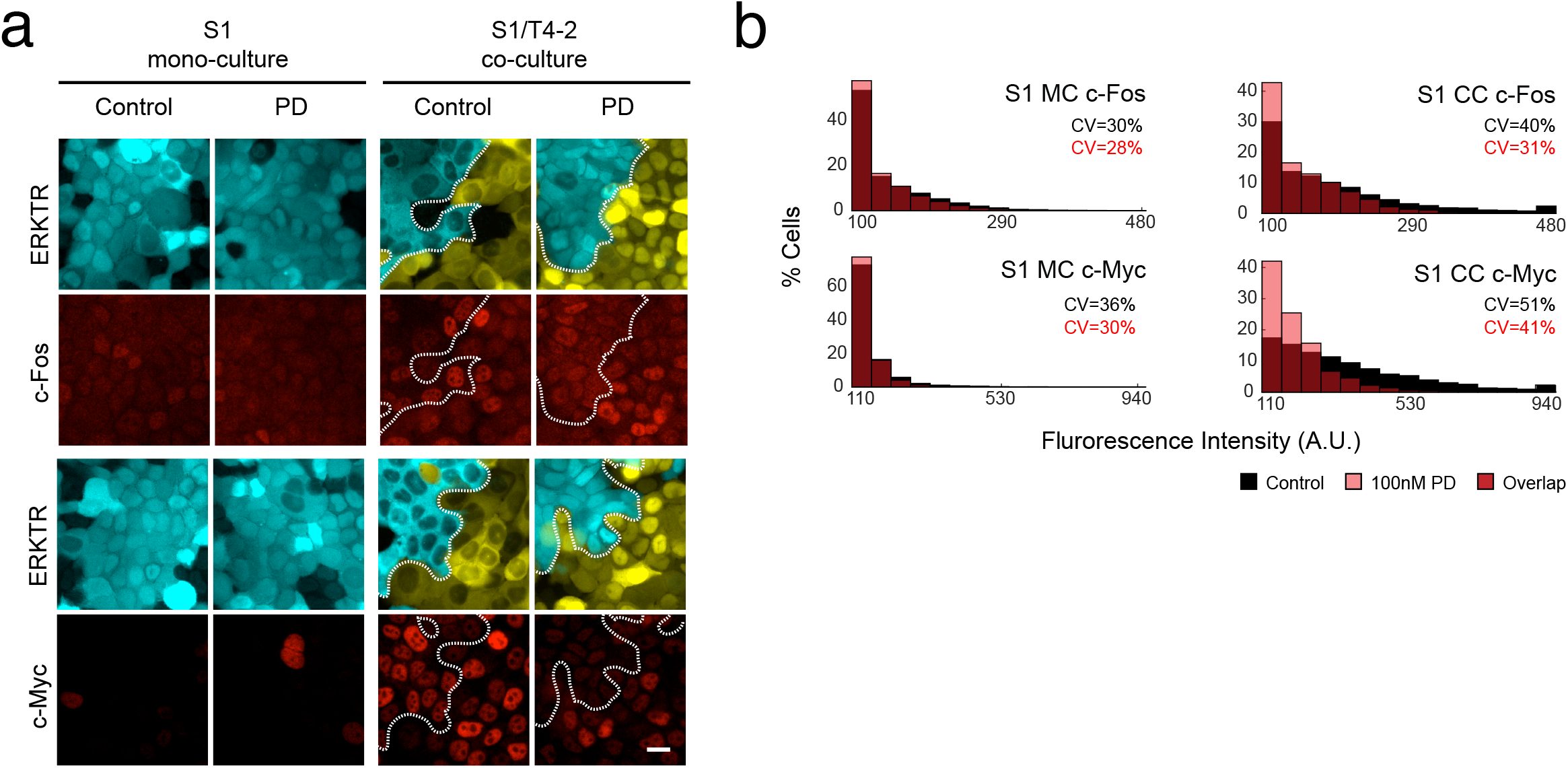
Heterogeneous expression of c-Fos and c-Myc driven by paracrine Areg. (A) Mono-cultured S1 (turquoise) and co-cultured S1 and T4-2 (yellow) cells stained for ETG expression of c-Fos and c-Myc (red) under control or PD conditions. Dashed lines indicate division between S1 and T4-2 cells in co-culture images. (B) Histogram of c-Fos and c-Myc protein expression distributions obtained from immunofluorescence staining. Control treated data is depicted in black, PD treated in pale red, and overlapping data in dark red.

**Supplemental Figure 6:**
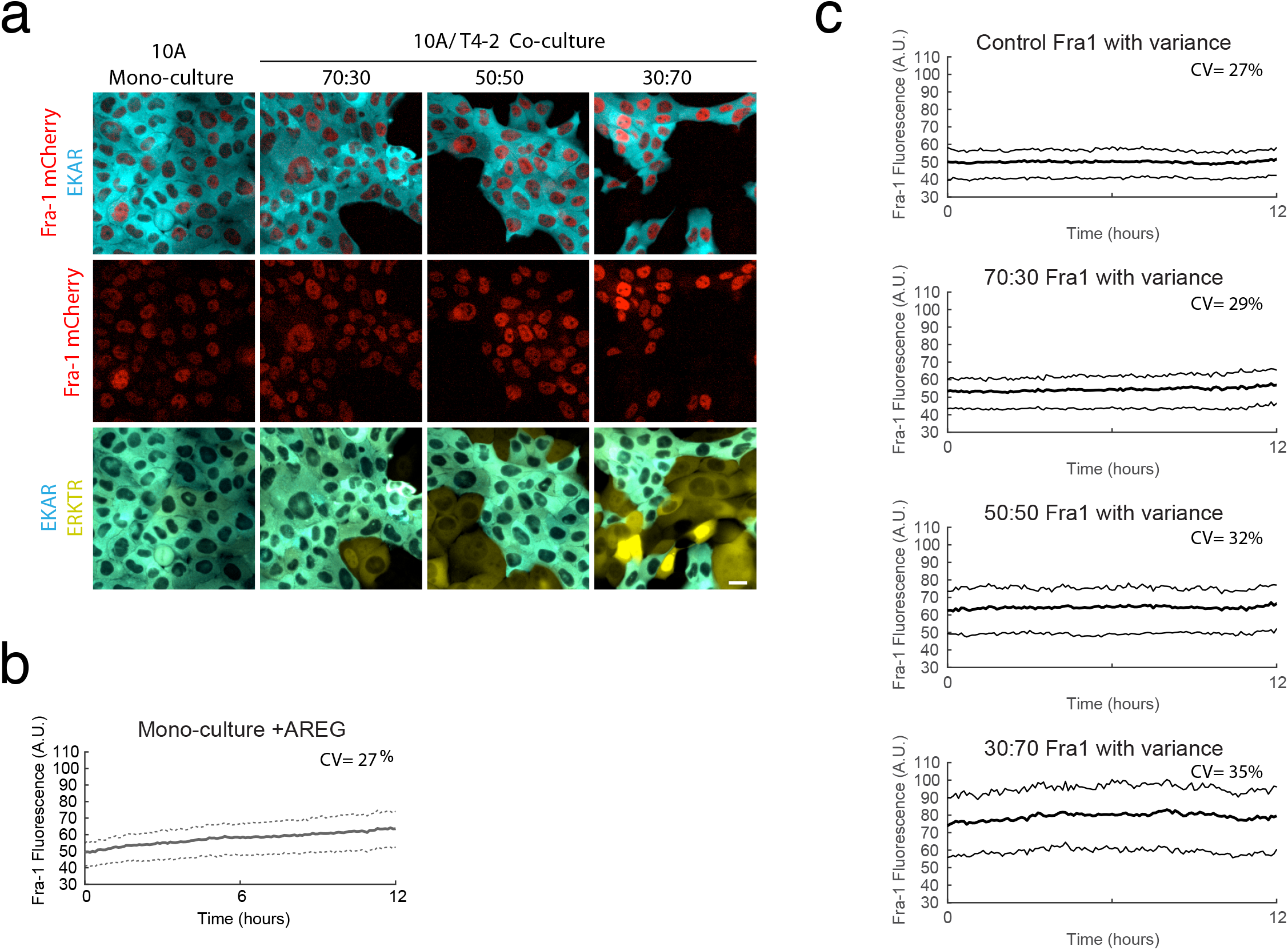
Real-time measurement of paracrine induced Fra-1 expression heterogeneity. (A) Still images if MCF10A cells carrying the EKAR reporter (turquoise) and mCherry tagged endogenous Fra-1 in mono-culture or co-culture with T4-2 cells carrying the ERKTR reporter (yellow). (B) Mean Fra-1::mCherry expression following the addition of 20ng/ml Areg in mono-culture conditions. Dashed line represent expression variance as the 25^th^ and 75^th^ interquartile range. (C) Mean Fra-1::mCherry expression under the indicated conditions. Dashed lines represent the variance as above.

**Supplemental Figure 7:**
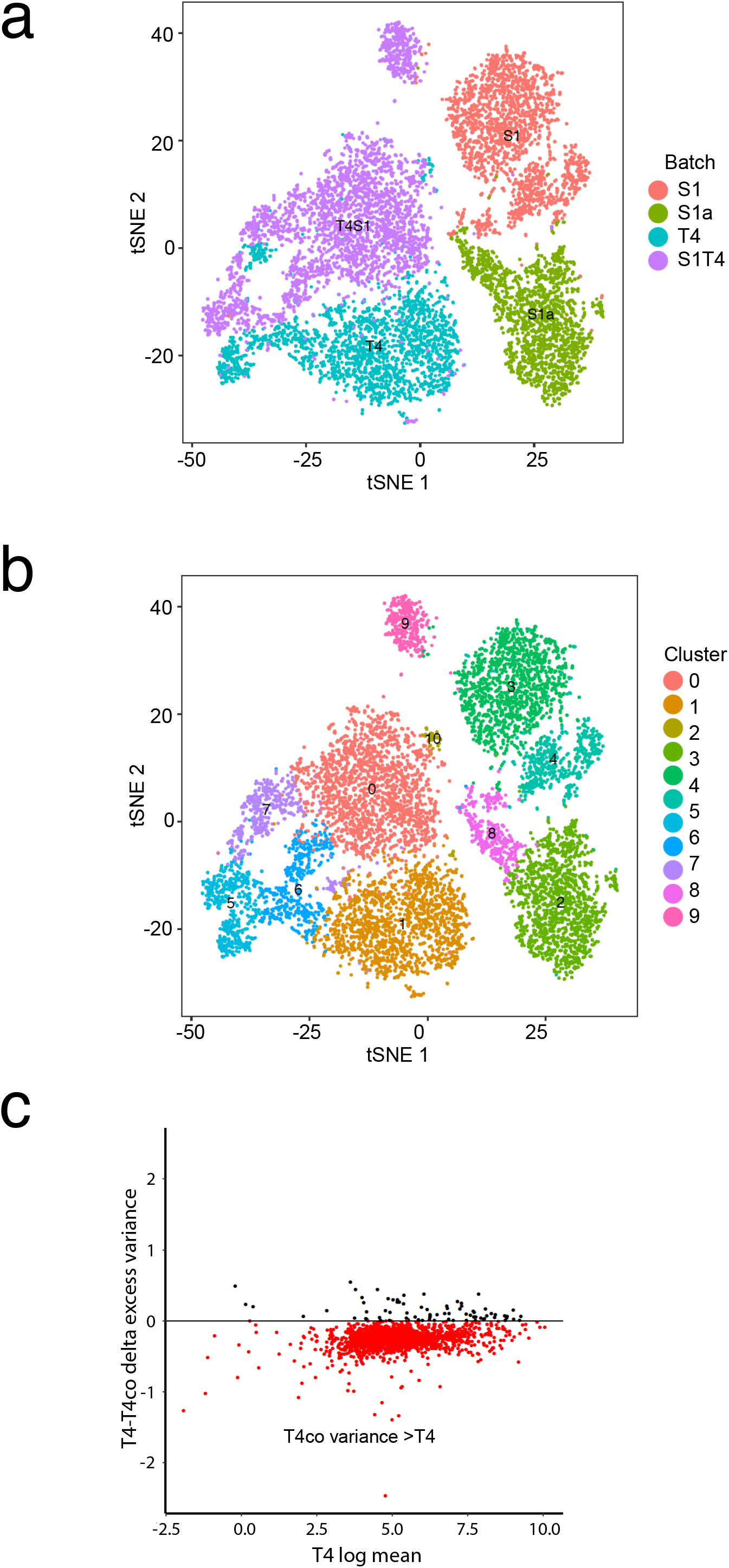
Single-cell transcriptomics and excess variance including T4-2 co-culture comparison. (A) tSNE plot of S1 control (orange), S1 treated with 100ng/ml amphiregulin (S1a, green), T4-2 control (blue), or S1/T4-2 co-cultured cells ((T4S1) purple), clustered based on single cell transcript profiles by cell type and color-labeled by origin. (B) tSNE plot from cluster analysis that included T4-2 co-cultured cells with 11 clusters representing distinct transcriptional profiles. (C) Difference in excess gene expression variance between T4-2 control cells (T4) and T4-2 co-cultures with S1 cells (T4co) plotted against the log mean T4 variance. Dots represent a single gene from ~1900 genes sampled. Red dots represent genes with displaying an increase in excess variance in T4co compared to T4. Black dots represent genes displaying a decreased in excess variance in T4co compared to T4.

**Supplemental Figure 8.**
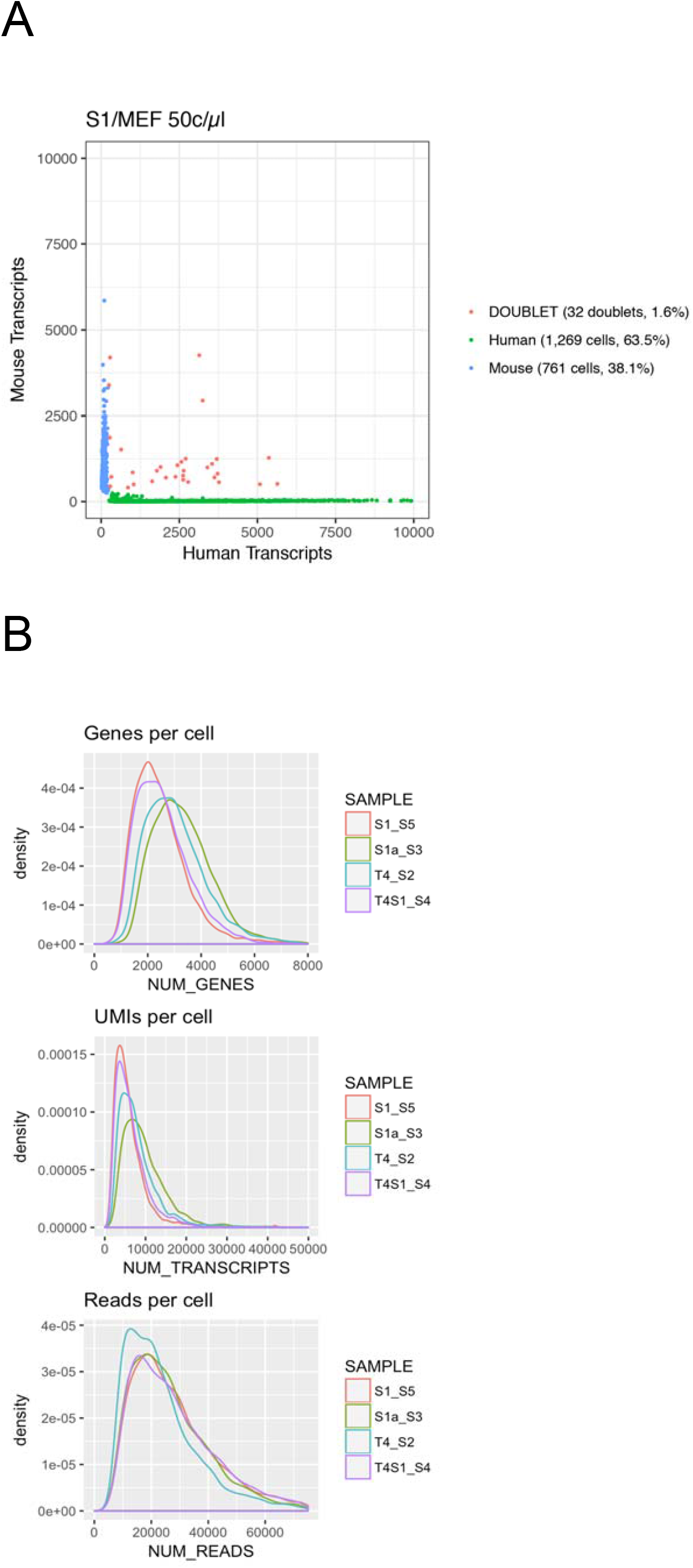
(A) Species mixing experiment. Mixtures of human (HEK) and mouse (MEF) cells were analyzed by Drop-seq at a concentration of 50c/µl (after droplet formation). The plot shows the number of human and mouse transcripts associating with each STAMP. Blue dots indicate STAMPs that were designated as mouse-specific; green dots indicate STAMPs that were human-specific and orange indicate STAMPS associated with a mixture of human and mouse transcripts. (B) DropseqStats. Number of genes, UMIs and reads per cell recovered across libraries.

### Supplemental Movies

**Supplemental Movie 1- Mono-cultured S1 and T4-2 ERK dynamics:** (A) Control (without growth factors) S1 cells compared to EGF stimulated S1 cells carrying the ERKTR reporter. (B) Control (without growth factors) T4-2 cells compared to EGF stimulated T4-2 cells carrying the ERKTR reporter. (C) Control (without growth factors) T4-2 cells compared to PD (MEK inhibitor) treated T4-2 cells carrying the ERKTR reporter. Time scale = hours.

**Supplemental Movie 2- S1/T4-2 ratio dependent ERK dynamics in co-culture:** (A) Mono-cultured S1-ERKTR mTurquoise cells. (B) Mono-cultured T4-2-ERKTR mVenus cells. (C) 10:90 S1 to T4-2 ratio co-culture. (D) 30:70 S1 to T4-2 ratio co-culture. (E) 50:50 S1 to T4-2 ratio co-culture. (F) 70:30 S1 to T4-2 ratio co-culture. (G) 90:10 S1 to T4-2 ratio co-culture. Time scale = hours.

**Supplemental Movie 3- Co-cultured S1 and T4-2 ERK dynamics:** (A) Control (without growth factors) S1/T4-2 co-cultured cells compared to EGF stimulated cells carrying the ERKTR reporter. (B) Control (without growth factors) S1/T4-2 co-cultured cells compared to erlotinib inhibited cells carrying the ERKTR reporter. Time scale = hours.

**Supplemental Movie 4- MCF10A-EKAR3-Fra-1::mCherry cells in mono and co-culture conditions:** (A) Mono-cultured MCF10A cells without exogenous stimulation. EKAR3 reporter left panel, Fra-1::mCherry right panel (red). (B) MCF10A cells co-cultured with T4-2 cells (unlabeled cells in field of view). EKAR3 reporter left panel, Fra-1::mCherry right panel (red). EKAR3 reporter white represents the ‘low’ ERK activity state, brownish-red the ‘high’ activity state. Time scale = hours.

**Supplemental Table 1**. Drop-seq. Cell concentrations in cell syringe, droplet sizes generated by the microfluidics device, effective cell concentrations, cell concentration in droplets, anticipated STAMPs processed per 50µl PCR reaction and number of cycles for each library.

**Supplemental Table 2**. Cells numbers and statistics for Drop-seq libraries after cell filtering

**Supplemental Table 1:** Single cell sequence parameters.

| Cells | Cell conc<br>loaded<br>[c/μl] | Droplet size<br>[nl] | Effective<br>cell conc<br>[c/μl] | In drop<br>[c/μl] | Stamps<br>per PCR<br>reaction | # cycles<br>PCR |
| --- | --- | --- | --- | --- | --- | --- |
| S1/MEF | 125 | 0.8 | 100 | 50 | 100 | 12 |
| T4 | 120 | 0.975 | 117 | 58.5 | 100 | 12 |
| T4/S1 | 120 | 0.975 | 117 | 58.5 | 100 | 15 |
| S1 | 100 | 0.975 | 97.5 | 48.75 | 100 | 15 |
| S1 amphiregulin | 125 | 0.8 | 100 | 50 | 100 | 14 |

**Supplemental Table 2:**
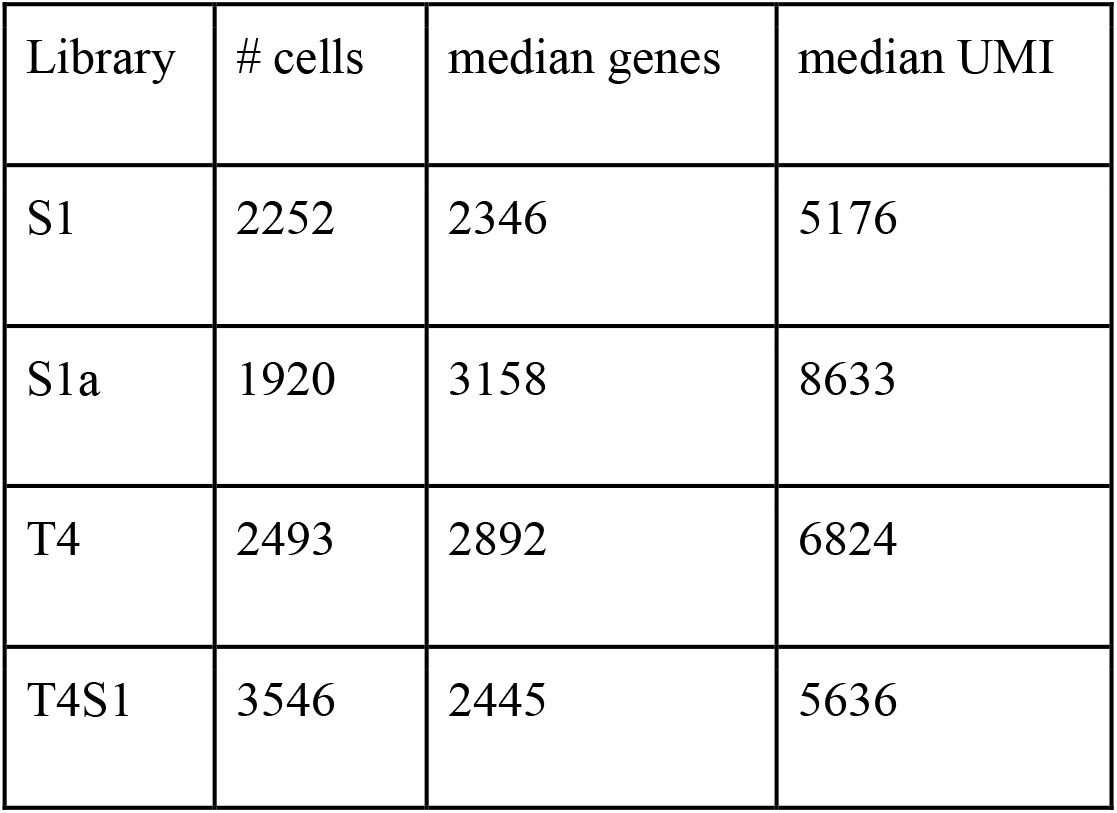
Cells numbers and statistics for Drop-seq libraries after cell filtering.

| Library | # cells | median genes | median UMI |
| --- | --- | --- | --- |
| S1 | 2252 | 2346 | 5176 |
| S1a | 1920 | 3158 | 8633 |
| T4 | 2493 | 2892 | 6824 |
| T4S1 | 3546 | 2445 | 5636 |

## Acknowledgements

Funding for this work was provided by the American Association for Cancer Research Stand Up To Cancer (SU2C-AACR-IRG-01-16) to J.G.A. Stand Up To Cancer is a program of the Entertainment Industry Foundation. Research grants are administered by the American Association for Cancer Research, the scientific partner of SU2C. A.E.D. was funded by the California Institute of Regenerative Medicine/Children’s Hospital of Oakland Clinical Research Fellowship and A.E.D. and M.J.B. were funded by the Breast Cancer Research Foundation and the Woodland Fund. We would like to thank the members of the Bissell and Albeck Labs for their helpful suggestions on the manuscript.

## Author Contributions

A.E.D. and J.G.A. conceptualized the study, analyzed data, and wrote the manuscript. A.E.D., S.J.T., and A.R.R. performed imaging experiments. A.E.D. and S.S. performed single cell RNA sequencing. T.E.G., S.S., M.P., G.Q. performed data processing and statistical analysis. M.J.B. and C.J. analyzed data and contributed to writing the manuscript.

The authors declare no competing interests

